# Programmable acetylation modification of bacterial proteins by a Cas12a-guided acetyltransferase

**DOI:** 10.1101/2022.07.29.502094

**Authors:** Yanqiang Liu, Ni Zuo, Weihong Jiang, Yang Gu

## Abstract

Protein lysine acetylation (PLA) is a crucial post-translational modification in organisms that regulates a variety of metabolic and physiological activities. Many advances have been made in PLA-related research; however, the quick and accurate identification of causal relationships between specific protein acetylation events and phenotypic outcomes at the proteome level remains unattainable due to the lack of in situ targeted modification techniques. In this study, based on the characteristics of transcription-translation coupling in bacteria, we designed and constructed an in situ targeted protein acetylation (TPA) system fusing the dCas12a protein, guiding element crRNA, and bacterial acetylase At2. Rapid identification of multiple independent protein acetylation and cell phenotypic analyses in gram-negative *E. coli* and gram-positive *C. ljungdahlii* demonstrated that TPA is a specific and efficient targeting tool for protein modification studies and engineering.

## INTRODUCTION

The post-translational dynamic covalent modification of proteins includes phosphorylation, acylation, methylation and other forms. Together with transcriptional and translational regulations, these modifications form a multi-level metabolic regulation network in organisms. This network achieves a precise control of various physiological and metabolic activities to cope with the changing external environment. Among them, protein lysine acetylation (PLA) is an important and widespread post-translational modification. It is a reversible dynamic process conserved in all life systems. According to the catalytic mechanism, PLA is divided into non-enzymatic and enzymatic catalysis; the former is a spontaneous chemical reaction using acetyl phosphate as the acetyl donor and the latter is catalysed by acetyltransferase with acetyl-CoA as the acetyl donor (1). PLA cannot only directly affect the structure and activity of proteins but also intervene in protein-protein or protein-nucleic acid interactions. It is involved in various aspects of organisms, such as metabolism, growth, and development (2-4). Furthermore, its abnormalities can cause metabolic disorders and diseases (5). Several studies have shown the universality and importance of PLA; however, the functions of the vast majority of PLA modifications remain unclear. In addition, it is impossible to conduct in situ modification intervention at the proteomic level, mainly due to the lack of efficient research tools for targeted modification.

Current methods adopted in PLA research mainly include: (i) perturbation of global acetylation levels by either overexpressing/inactivating genes encoding acetyltransferase, deacetylase, and acetylphosphate metabolism-related enzymes (6,7) or using chemical inhibitors (8,9); (ii) acetylation and deacetylation-mimicking mutations by changing lysine residues (K) to glutamine (Q) and arginine (R), respectively (10-12). This method cannot simulate the real acetylation status of the target protein, and may alter the amino acid sequence, possibly affecting protein structure and function; (iii) site-specific replacement of the target lysine residue with acetyllysine by introducing an amber codon UAG into the corresponding gene, a pyrrolysyl-tRNA synthetase/tRNA_CUA_ pair, and an exogenous acetyllysine (13). This is suitable for functional verification of the acetylation on a single lysine residue of interest. Collectively, these methods have their own characteristics and are helpful in PLA research, but none of them can achieve targeted acetylation at the proteomic level to rapidly reveal causal relationships between specific protein modifications and phenotypes. Therefore, new approaches must be developed.

Bacteria play a crucial role in the fields of medical health, environmental protection and biological manufacturing, and their protein modification has attracted extensive attention. A significant feature of bacteria is the coupling of transcription and translation. Based on this concept, we investigated the use of CRISPR-Cas tools to guide an acetyltransferase towards target genes, thereby enabling specific acetylation of their encoding proteins (Fig 1A). To this end, we fused the nuclease-inactivated dCas12a with suitable acetylases, designed and constructed an in situ targeted modification system for targeted protein acetylation (TPA), and characterised its efficiency in both gram-positive and gram-negative bacteria. Using TPA, we achieved not only specific in vivo acetylation of multiple crucial proteins without detectable off-target acetylation events, revealing the consequent phenotypic changes, but also rapid identification of an acetylation-regulated sigma factor by screening a mini TPA library. Moreover, the TPA-like target protein modification was further extended to propionylation, demonstrating the potential universality of this tool.

**Fig 1.**
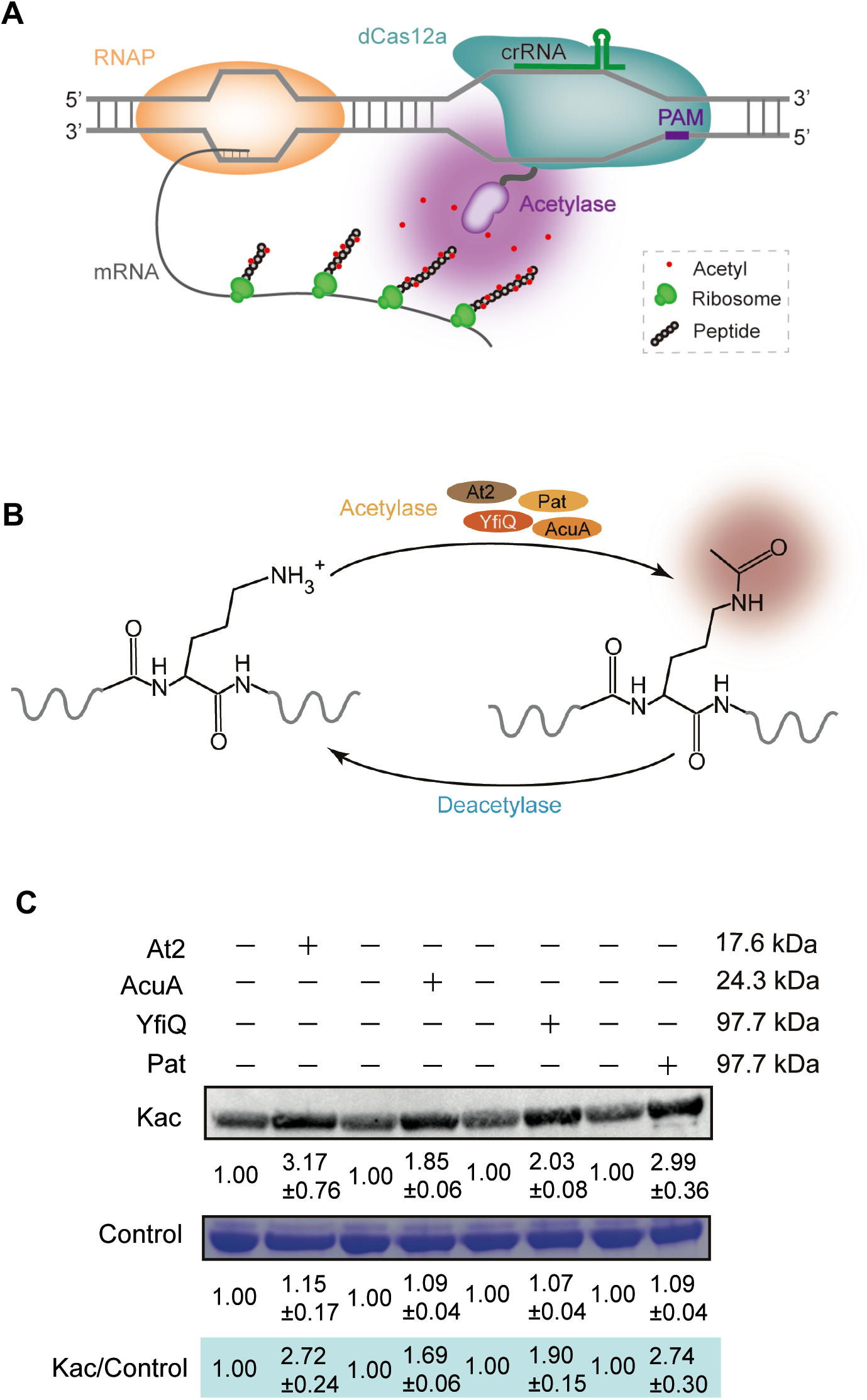
Design of the in vivo target protein acetylation (TPA) system in bacteria. (**A**) The proposed action mechanism of TPA in bacteria. A fusion protein containing a dCas12a and acetyltransferase proteins mediates the crRNA-guided specific acetylation of target proteins based on the synchrony of gene transcription and protein translation in bacteria. (**B**) Reversible protein lysine acetylation mediated by acetyltransferases and deacetylases. (**C**) Screening of an appropriate acetyltransferase for the dCas12a-acetylase fusion. The candidates include *Salmonella enterica* Pat, *Escherichia coli* YfiQ, *Bacillus subtilis* AcuA, and *Clostridium ljungdahlii* At2. Western blot analysis was performed to examine the in vitro acetylation activities of these fusion proteins towards the *C. ljungdahlii* Pta protein. Kac, lysine (K) acetylation. The data are presented as mean ± SD from three independent experiments.

## MATERIAL AND METHODS

### Bacteria, media, reagents, and strain cultures

The *E. coli* DH5α and BL21 (DE3) as well as their derived strains were grown in the lysogeny broth (LB) medium. The *E. coli* BW25113 strain was grown in LB medium or standard minimal medium (14) supplemented with 10 mM acetate. Chloramphenicol (12.5 μg mL^−1^), kanamycin (100 μg mL^−1^), and nicotinamide (an inhibitor of deacetylase, 0.5 mM) were added to the media when needed. The *C. ljungdahlii* strain (DSM 13528) was first grown in YTF medium (15) for inoculum preparation, which was then transferred into a modified ATCC medium 1754 (2-fold increased concentration of all the mineral elements; fructose was removed) with a headspace of CO−CO_2_−H_2_−N_2_ (56%, 20%, 9%, and 15%; pressurised to 0.2 MPa) for gas fermentation, in which 5 μg mL^−1^ of thiamphenicol was added for plasmid selection. Anaerobic cultivation was performed in an anaerobic chamber (Whitley A35 Anaerobic Workstation, Don Whitley Scientific Limited, Bingley, West Yorkshire, UK).

Chemical reagents were purchased from Sigma-Aldrich (St Louis, USA) and Aladdin (Shanghai Aladdin Biochemical Technology Co., Ltd., Shanghai, China). KOD plus Neo and KOD FX DNA polymerase (Toyobo, Osaka, Japan) were used for PCR amplification. The assembly of multiple DNA segments for plasmid construction was performed using the ClonExpress MultiS One Step Cloning Kit (Vazyme Biotech Co., Ltd., Nanjing, China). The restriction enzymes and ligase used in plasmid construction were purchased from Thermo Fisher Scientific (Thermo Fisher Scientific, Vilnius, Lithuania) and Takara (Takara, Dalian, China), respectively. Plasmid isolation and DNA purification were performed using kits (Axygen Biotechnology Company Limited, Hangzhou, China). All the primers used in this study were synthesised by GenScript (GenScript, Nanjing, China) and BioSune (Biosune, Shanghai, China).

### Plasmid construction

All the primers and plasmids used in this study are shown in Supplementary Tables S1 and S2, respectively.

The plasmid pNat2-L2-Dc-acs-NTs that carried *At2* and the *Francisella tularensis dcas12a*, together with a crRNA expression cassette, was constructed as follows: the plasmid pACYC-Duet1 was treated with NdeI and XhoI; next, the *At2* fragment was cloned through PCR amplification using the *C. ljungdahlii* genomic DNA as the template and the primers At2-F/At2-L2-R; the *dCas12a* fragment was cloned through PCR amplification using the pUC57-dCas12a as the template and the primers dCas12a-L2-F/dCas12a-R, and then linked to the the At2 fragment by overlapping PCR using the primers At2-F/dCas12a-R, yielding the DNA fragment At2-L2-dCas12a; finally, the At2-L2-dCas12a fragment and the aforementioned linear pACYC-Duet1 plasmid were assembled using a ClonExpress II one-step cloning kit (Vazyme Biotech Co., Ltd., Nanjing, China), yielding the plasmid pNat2-L2-Dc-NCr; then, plasmid pNat2-L2-Dc-NCr was treated with NcoI and BamHI; next, the acs-NTs fragment was cloned through overlapping PCR using the primers acs-NTs-F/acs-NTs-R; the acs-NTs-L fragment was cloned through PCR amplification using the acs-NTs as the template and the primers CrRNA-F/CrRNA-R; finally, the acs-R-L fragment and the aforementioned linear pNat2-L2-Dc-NCr plasmid were assembled using a ClonExpress II one-step cloning kit (Vazyme Biotech Co., Ltd., Nanjing, China), yielding the plasmid pNat2-L2-Dc-acs-NTs.

The other plasmids for targeted acetylation in *E. coli*, including pNat2-L2-Dc-acs-Ts and pNat2-L2-Dc-rnb-NTs, targeting the template strand of *acs* and non-template strand of *rnb*, respectively, were constructed in a similar manner, except that the crRNA expression cassettes were changed accordingly.

The plasmid pET28a-pta for *pta* expression (for the following protein purification) was constructed as follows: the *pta* gene was amplified by PCR using the *C. ljungdahlii* genomic DNA as the template and the primers pET28a-Pta-F/pET28a-Pta-R; then, the *pta* fragment and the linear plasmid pET28a, which was obtained by double digestion with NdeI and XhoI, were assembled using the ClonExpress II one-step cloning kit, yielding the target plasmid pET28a-Pta.

The other plasmids for expressing *yfiQ, pat, acuA, at2, Ndc-L1-At2, Ndc-L2-At2, Ndc-L3-At2, Ndc-L4-At2, Nat2-L2-Dc*, and *acs* were constructed in a similar manner, except that gene sequences were changed accordingly.

The plasmid pTargetF-N20-acs that was used to insert a 6×His tag (HHHHHH) at the C-terminus of the *acs* gene on the *E. coli* chromosome was constructed by PCR amplification using the plasmid pTargetF as the template and the primers acs-6H-F/acs-6H-R.

The other plasmids for inserting a 6×His tag at the C-terminus of the *rnb* genes were constructed in a similar manner, except that the crRNA was replaced accordingly.

### Gene expression and protein production

Purification of the Hig-tagged proteins encoded by *pta, yfiQ, pat, acuA, at2, Ndc-L1-At2, Ndc-L2-At2, Ndc-L3-At2, Ndc-L4-At2, Nat2-L2-Dc*, and *acs* was performed as previously described (16). Briefly, gene expression was induced at 16°C overnight by adding 0.5 mM Isopropyl-β-D-thiogalactoside (IPTG) when A_600_ reached to 0.6–0.8. Cells were harvested by centrifugation (6 min, 5,000 ×g, 4°C), and then disrupted by a French Press (Constant Systems Limited, UK). Cell debris was separated from the soluble fraction by centrifugation (30 min, 20,000 g, 4°C). The soluble fraction was loaded onto a Ni Sepharose™ 6 Fast Flow agarose (GE Healthcare) column for protein purification. Proteins were eluted using a buffer (pH 7.9) containing 20 mM Tris–HCl, 200 mM KCl, 10% (v/v) glycerol, and 500 mM imidazole. The eluent fraction was then transferred to an Amicon Ultra-15 Centrifugal Filter (Millipore, Billerica, MA) and eluted for three times by using a buffer (pH 7.9) containing 20 mM Tris–HCl, 500 mM KCl, and 10% (v/v) glycerol. The purified proteins were stored at −80°C. All the proteins purified in this study are shown in Supplementary Figure S1.

A 6×His tag was inserted at the C-terminal of *rpoC* on the chromosome (Supplementary Figure S2), aiming at the following RNAP holoenzyme purification.

Additionally, a 3×flag tag was inserted at the C-terminal of *rpoD* on the chromosome (Supplementary Figure S2), aiming at the following detection of RpoD in cell lysates by western blot analysis using an anti-flag antibody. The RNAP holoenzyme purification procedure was the same as reported previously (17).

### RNA purification and real-time quantitative PCR (qPCR)

*E. coli* cells were grown at 37°C in LB or standard minimal medium supplemented with 10 mM acetate with shaking at 220 rpm for 3–12 h. Cells were harvested by centrifuging the culture at 5, 000 *g* for 10 min at 4 °C. Total RNA was then isolated using a kit (catalogue no. cw0581; CWBIO) according to the manufacturer’s protocol, and further treated with DNase I (TaKaRa) to eliminate residual DNA contained in the extracted samples. RNA concentration was determined using a NanoDrop spectrophotometer (Thermo Fisher Scientific, Waltham, MA, USA). Complementary DNA (cDNA) was obtained by reverse transcription using a PrimeScript RT reagent kit (TaKaRa) according to the manufacturer’s instructions. RT-qPCR was performed as described previously (18). All the reactions were run in technical triplicate. The *gapA* (NP_416293.1) gene was used as an internal control.

*C. ljungdahlii* cells were cultured anaerobically at 37°C in modified ATCC medium 1754 (19) with a headspace of CO−CO_2_−H_2_−N_2_ mixture (56%, 20%, 9%, and 15%; pressurised to 0.2 MPa). Cells were harvested when they reached an optical density (OD_600_) of 1.0. The following RNA purification and RT-qPCR were performed as previously described for *E. coli*.

### Western blotting

Bacterial cells were washed twice with protein purification buffer and disrupted by a French Press (Constant Systems Limited, UK). Cell debris was separated from the soluble fraction by centrifugation (20 min, 20,000 g, 4°C). Protein concentrations were determined using a kit (Sangon Biotech Co., Ltd., China). Next, the samples were loaded onto a 12% SDS-PAGE gel for protein separation by electrophoresis, and the gel was then transferred to a nitrocellulose membrane (GE Healthcare) for 20–50 min at 400 mA. The membrane was blocked with 1×Tris-buffered saline and Tween 20 (TBST: 10 mM Tris-HCl, pH 7.4, 100 mM NaCl and 0.2% Tween 20) containing 5% non-fat dry milk for 20 min. Membranes were then stained with anti-acetyllysine antibody or anti-Propionyllysine antibody (the primary antibody), diluted in TBST–0.5% non-fat dry milk (1.5: 20,000) for 1 h. The blots were washed with TBST (TBS + 0.5% Tween 20) three times, and then incubated with goat horseradish peroxidase-conjugated anti-rabbit or anti-mouse antibody (the secondary antibody) that was diluted in TBST (1: 5,000) for 1 h, followed by three washes with TBST. Finally, the membranes were imaged using an enhanced chemiluminescence (ECL) system (ImageQuant LAS 4000 Mini; GE) according to the manufacturer’s instructions.

### In vitro acetylation assay

Acetylation activity assays were performed as previously described (20). Briefly, a 60 μL reaction system containing 50 mM Tris-HCl (pH 8.0), 2 μM target proteins, 0.5 μM acetyltransferase, 0.2 mM acetyl-CoA, and 5% glycerol was constructed and incubated at 37°C for 3 h. After the reaction, the samples were used for western blot analysis, yielding relative acetyltransferase activity.

#### Mass spectrometry measurements

Identification of acetylated lysine residues was achieved by mass spectrometry. The purification procedure of the proteins acetylated in vivo was the same as previously described (16). The purified proteins were separated by 12% SDS-PAGE. The excised bands containing the target proteins were digested with trypsin at 37°C for 20 h. The obtained peptides were further separated by a nano-liquid chromatography system (Easy-nLCII, Thermo Fisher Scientific) (mobile phase solvents A [0.1% (vol/vol) formic acid in water] and B [0.1% (vol/vol) formic acid in 84% (vol/vol) acetonitrile]) and then analyzed with a Q-Exactive mass spectrometer. The mass spectrometer was operated as previously described (21).

### Gas fermentation

Gas fermentation of the wild-type *C. ljungdahlii* strain and its derivatives was carried out in 125 mL serum bottles (Sigma-Aldrich, Merck Life Science, Gillingham, UK) with a working volume of 30 mL. For inoculum preparation, 200 μL of frozen stock was inoculated into 5 mL of the YTF medium (15) and incubated anaerobically at 37°C for 24 h. When the optical density (OD_600_) of the cells reached 0.8–1.0, 5% (v/v) of the inoculum was transferred to 30 mL of the ATCC medium 1754 with a headspace of a CO/CO_2_/H_2_/N_2_ mixture (56%, 20%, 9%, and 15%; pressurised to 0.2 MPa) for gas fermentation. The gases in the headspace were replaced every 24 h. Media preparation and strain inoculation were performed using a Whitley A35 anaerobic workstation (Don Whitley Scientific, West Yorkshire, UK).

### Analytical methods

Cell growth was measured based on the absorbance of the culture at A600 (OD_600_) using a spectrophotometer (DU730; Beckman Coulter). The fermentation products were assayed as previously reported (19). Briefly, bacterial cultures were collected and centrifuged at 7,000 × *g* for 10 min at 4°C. The supernatant was used for the assays of acetate and ethanol assays. The concentrations of acetate and ethanol were determined using a 7890A gas chromatograph (Agilent, Wilmington, DE, USA) equipped with a flame ionisation detector (Agilent) and a 3.0 m × 0.32 mm capillary column (EC-Wax, Alltech, Lexington, KY) packed with fused silica. The analysis was performed as follows: oven temperature, programmed from 85 to 150°C at a rate of 25°C/min; injector temperature, 250°C; detector temperature, 300°C; nitrogen (carrier gas) flow rate, 20 ml/min; hydrogen flow rate, 30 ml/min; air flow rate, 400 ml/min; and internal standaeds, isobutanol, isobutyric acid, and hydrochloric acid (for acidification).

## RESULTS

### Selection and characterisation of acetyltransferases for TPA editor design

An acetyltransferase with strong catalytic activity and small molecular weight fused to dCas12a would enable the resulting TPA construct to achieve high protein acetylation efficiency and low steric hindrance in DNA binding. Here, we chose four reported bacterial acetyltransferases, *Salmonella enterica* Pat (97.7 kDa), *Escherichia coli* YfiQ (97.7 kDa), *Bacillus subtilis* AcuA (24.3 kDa), and *Clostridium ljungdahlii* At2 (17.6 kDa) (22-25) for *in vitro* characterisation (Fig 1B).

Pta, a recently reported acetylated protein in *C. ljungdahlii* (25), was adopted as the target for treatment in vitro by the purified Pat, YfiQ, AcuA, and At2. Obviously increased acetylation levels of Pta were observed following treatment with all four acetyltransferases (Fig 1C), indicating their high catalytic activities. Considering the smallest molecular weight of At2 among the four tested enzymes, we selected it for the subsequent construction of the TPA editor.

### Construction of the TPA editor

We first constructed a fusion protein containing At2 and a dCas12a protein (DNase-deactivated Cas12a) from *Francisella tularensis*. In this design, At2 was fused to the C-terminus of dCas12a via a peptide linker (Fig 2A). Considering that linkers can have unfavourable impacts on fused proteins, we tested different peptide linkers in the construction of the dCas12a-At2 fusion, including short flexible (GGGS) (L1), short rigid (GSGEAAAK) (L2), medium rigid (GSG (EAAAK)_2_) (L3), and long rigid (GSG (EAAAK)_3_) linkers (L4) (26). Then, we assayed the acetyltransferase activity of the resulting four dCas12a-At2 fusions using the aforementioned At2-targeted protein, Pta, as the substrate (Fig 2A). Interestingly, compared with the control (free At2), the L2 linker gave the highest acetyltransferase activity to its corresponding fusion protein (Fig 2A), probably due to the increased stability and activity of the fused protein with a suitable peptide linker (27,28). Thus, the results suggested that L2 is a preferable linker for constructing the TPA editor. Furthermore, we fused At2 to the N-terminus of dCas12a using the L2 linker, yielding the At2-dCas12a fusion. As shown in Supplementary Figure S3, the At2-dCas12a and abovementioned dCas12a-At2 fusion proteins exhibited similar acetyltransferase activities, regardless of the relative location of At2 and dCas12a in the fusion. Thus, we adopted the At2-L2-dCas12a (abbreviated as At2-dCas12a) fusion for the following experiments.

**Fig 2.**
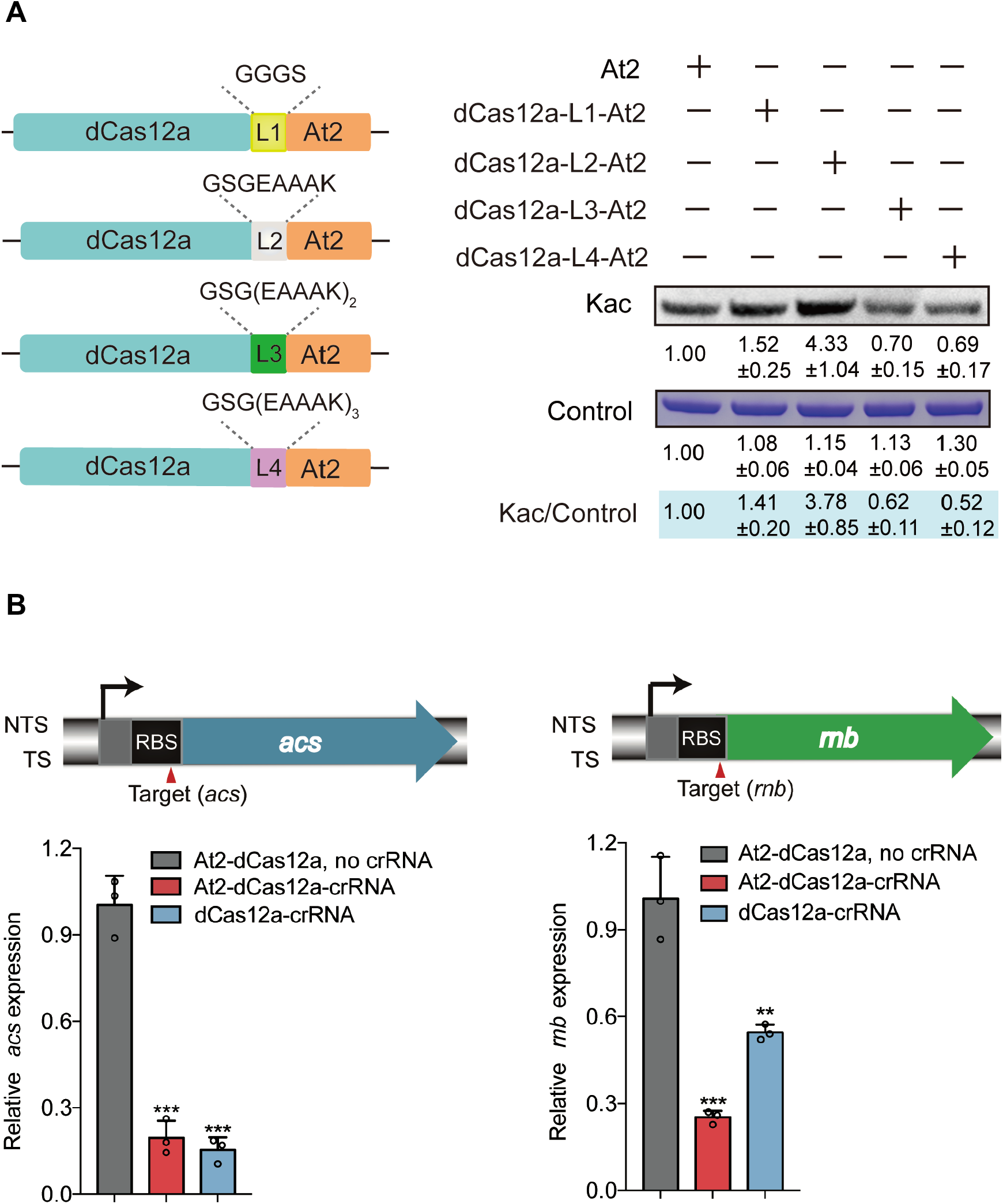
Construction of the TPA editor. (**A**) Screening of suitable peptide linkers for the dCas12a-acetyltransferase fusion. L1, GGGS; L2, GSGEAAAK; L3, GSG (EAAAK)_2_; L4, GSG (EAAAK)_3_. The plasmid-carried His-tagged fusion proteins were expressed in *Escherichia coli* and then purified for *in vitro* acetylation assays. Kac, lysine (K) acetylation. The data are presented as mean ± SD from three independent experiments. (**B**) DNA-binding activity of the At2-dCas12a fusion protein in *E. coli*. Two *E. coli* genes, *acs* (NP_418493.1) and *rnb* (NP_415802.1), were used as the targets for testing. RBS, ribosome binding site. Data are represented as mean ± SD (*n* = 3). Error bars show SDs. Statistical analysis was performed by a two-tailed Student’s *t*-test. **, *P* < 0.01 ***, *P* < 0.001, versus the control (At2-dcas12a, no crRNA).

Next, we investigated whether the At2-dCas12a fusion protein can specifically bind to target DNA in *E. coli* with the assistance of crRNAs. The *acs* and *rnb* genes, encoding acetyl-CoA synthetase and ribonuclease II in *E. coli*, were chosen for the test, and crRNAs targeting the RBS region of *acs* or *rnb* were designed (Fig 2B). Then, a plasmid carrying the At2-dCas12a fusion protein and the crRNA were transferred into *E. coli*; simultaneously, a plasmid expressing only the At2-dCas12a fusion protein (no crRNA) and a plasmid expressing dCas12a and crRNA were constructed as the negative and positive controls, respectively. As expected, both the plasmid containing all the elements and the positive control plasmid efficiently repressed the expression of target genes (Fig 2B), suggesting that the DNA-binding activity of dCas12a was not impaired by the fused At2.

### Validation of plasmid-carried TPA editors in gram-negative bacteria

Based on the above findings, we further evaluated the in vivo function of TPA. A plasmid (pNat2-L2-Dc-acs-Ts or pNat2-L2-Dc-acs-NTs) expressing the At2-dCas12a fusion protein and the corresponding crRNA was introduced into *E. coli* (Fig 3A), followed by transfer of a second plasmid (pET-28a-acs) encoding the His-tagged Acs protein (Fig 3A and Supplementary Table S1). To avoid the potential transcriptional repression of the At2-dCas12a fusion on *acs*, crRNAs were designed to target the position adjacent to the terminator codon (170 or 109 bp 5’ to the termination codon) (Fig 3B). The purified Acs protein was then used for western blot analysis. Encouragingly, we observed substantial acetylation of Acs in the presence of the At2-dCas12a fusion protein, while no such a phenomenon was detected in the control (At2-dCas12a, no crRNA) (Fig 3B), indicating that the designed TPA construct works well in *E. coli*. We further determined the acetylated lysine residues of Acs with and without TPA in vivo by mass spectrometry measurements, generating 12 and 15 sites, respectively, which were almost same except 131K, 161K, 226K, 541K, and 617K (Supplementary Table S3). Overall, these results suggested that the TPA editor could increase the acetylation level of Acs without causing great changes in acetylated sites compared to the normal condition (lacking TPA)

**Fig 3.**
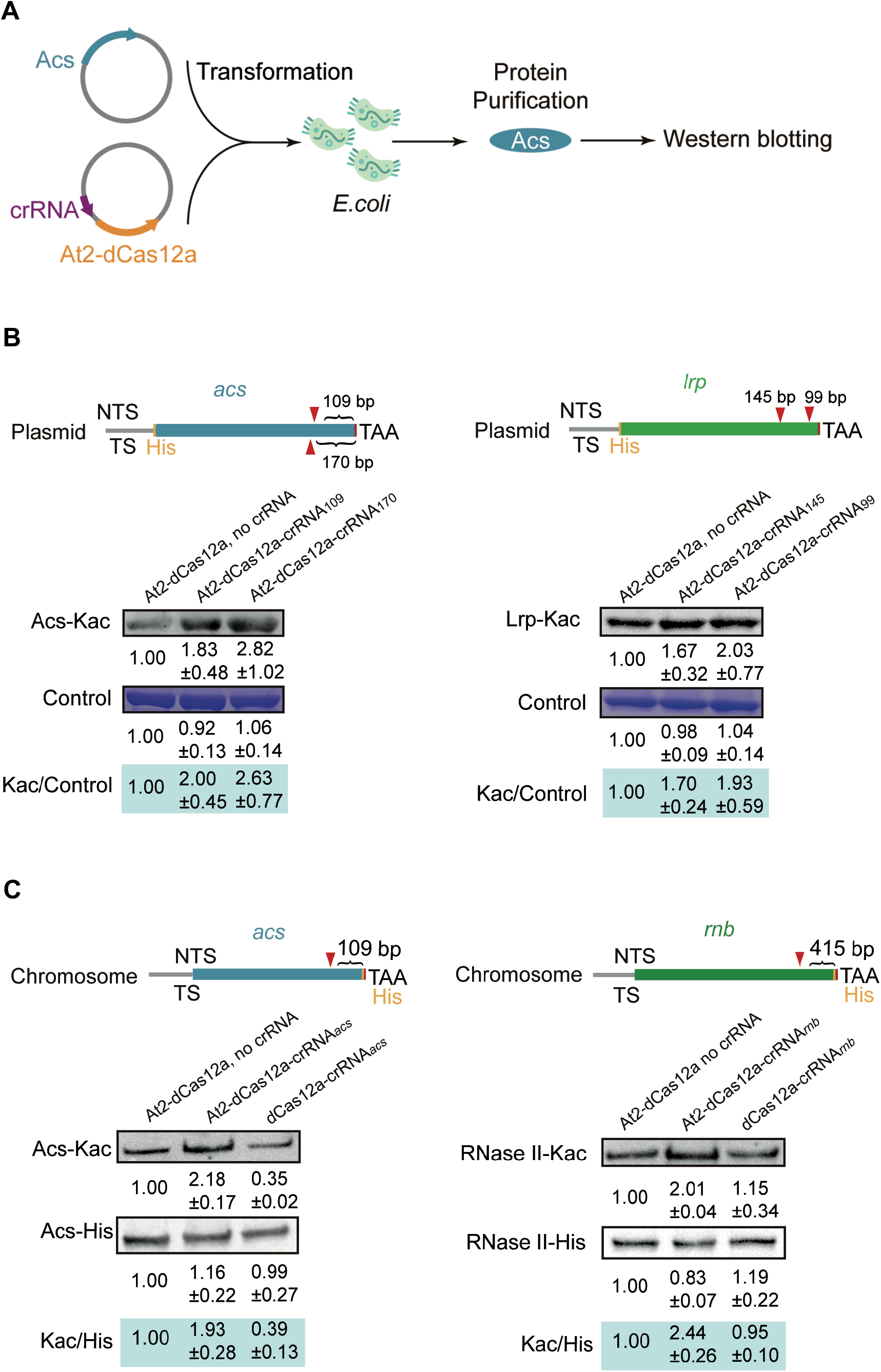
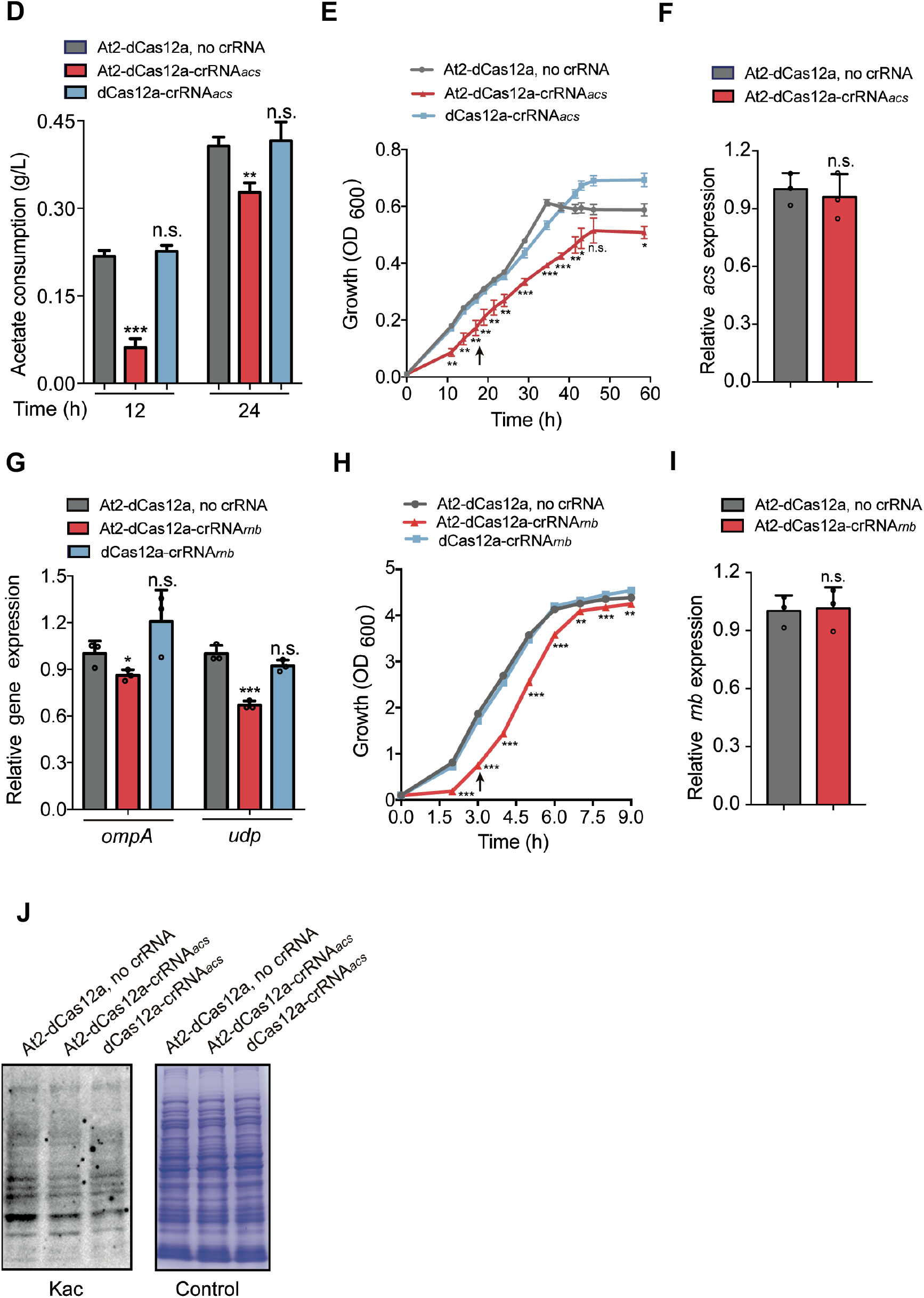
Validation of the TPA editor in *E. coli*. (**A**) Plasmids carrying the At2-L2-dCas12a fusion and the acs gene were transferred into *E. coli* to test the TPA effect. After targeted acetylation, His-tagged Acs was purified for western blot analysis. (**B**) Upper: TPA binding to the plasmid-carried *acs* and *lrp*. Bottom: the acetylation levels of Acs and Lrp with and without TPA editor. The control was the At2-dCas12a fusion (without crRNA). The data are presented as mean ± SD from three independent experiments. (**C**) Left: TPA binding to *acs* at the chromosome and acetylation levels of Acs with and without TPA editor. Right: TPA binding to *rnb* at chromosome and acetylation levels of RNase II with and without TPA editor. Cells were grown at 37°C in LB medium. The data are presented as mean ± SD from three independent experiments. (**D**) The acetate consumption of *E. coli* cells harboring the TPA editor (At2-dCas12a-crRNA_*acs*_), At2-dCas12a (no crRNA), and dCas12a-crRNA_*acs*_. Data are represented as mean ± SD (*n* = 3). (**E**) The growth of *E. coli* cells transferred with the TPA editor (crRNA_*acs*_), At2-dCas12a fusion (no crRNA), and dCas12a (with crRNA). (**F**) The transcriptional levels of *acs* in the *E. coli* cells transferred with the TPA editor and At2-dCas12a fusion (no crRNA). Data are represented as mean ± SD (*n* = 3). Cells were grown at 37°C in standard minimal medium supplemented with 10 mM acetate. The arrow represents the time point of cells collection for RT-qPCR. (**G**) The mRNA levels of *ompA* and *udp* in the *E. coli* cells harboring the TPA editor (At2-dCas12a-crRNA_*rnb*_), At2-dCas12a (no crRNA), and dCas12a-crRNA_*rnb*_. Data are represented as mean ± SD (*n* = 3) (**H**) The growth of *E. coli* cells transferred with the TPA editor (crRNA_*rnb*_), At2-dCas12a fusion (no crRNA), and dCas12a (with crRNA). (**I**) The transcriptional level of *rnb* in the *E. coli* cells transferred with the TPA editor and At2-dCas12a fusion (no crRNA). Cells were grown at 37°C in LB medium. Data are represented as mean ± SD (*n* = 3). (**J**) Global protein acetylation levels in the *E. coli* strains containing an active TPA editor (targeting *acs*), an untargeted TPA editor (no crRNA), and a single dCas12a-crRNA construct (control). ***, *P* < 0.001, **, *P* < 0.01, *, *P* < 0.05, n.s., no significance.

Such a dual-plasmid-based TPA system was further introduced into *E. coli* to target another gene, *lrp*, by replacing the original crRNA with new ones (145 and 99 bp 5’ to the termination codon of *lrp*) (pNat2-L2-Dc-lrp-Ts1 or pNat2-L2-Dc-lrp-Ts2; pET-28a-lrp) (Fig 3B and Supplementary Table S1). Lrp is a leucine-responsive global regulator and known to be acetylated in *E. coli* (29,30). After the in vivo TPA treatment, the Lrp protein was purified for western blot analysis. As expected, the acetylation levels of Lrp treated with the two At2-dCas12a-crRNA fusions were both obviously higher than that of control (At2-dCas12a, no crRNA). This finding further suggests that the TPA editor is functional in *E. coli*.

### TPA-based rapid identification of crucial acetylation events in *E. coli*

The real potential of TPA resides in its capacity to rapidly screen acetylation events capable of causing phenotypic changes of bacteria. To test this possibility, we focused on proteins associated with crucial physiological and metabolic processes in *E. coli*. The aforementioned Acs enzyme is known to be crucial for the conversion of acetate to acetyl-CoA in *E. coli*, and moreover, exhibits decreased activity after in vitro acetyltransferase treatment (3); however, whether the altered acetylation levels of Acs causes phenotypic changes remains unclear. Therefore, we first inserted a 6×His tag sequence at the chromosomal C-terminus of the *acs* gene in *E. coli* (Supplementary Figure S2); then, an At2-dCas12a-crRNA-expressing plasmid and two control plasmids were separately introduced into this *E. coli* (Supplementary Figure S4), yielding three strains for cultivation. Purified His-tagged Acs proteins from cell lysates were used for western blot analysis. The acetylation level of Acs was obviously increased in the *E. coli* strain containing the At2-dCas12a-crRNA-expressing plasmid (Fig 3C). This indicates that TPA can target to *acs*, enabling site-specific acetylation of the *acs*-encoding protein (Acs). Based on this finding, another crucial protein, RNase II, which is known to be acetylated in *E. coli* (31), was used as a target to further validate TPA (Supplementary Figure S2). As expected, the acetylation level of this protein was also obviously increased in the presence of the At2-dCas12a-crRNA construct (Fig 3C).

Next, we examined influences of targeted acetylation of Acs and RNase II on *E. coli*. For Acs, it has been known that this enzyme could catalyze the synthesis of acetyl-CoA from acetate and the increased acetylation level of Acs could block this process (3). Therefore, the interference of TPA on the acetylation of Acs would affect the consumption of acetate in *E. coli*. As expected, the fermentation results showed that the introduction of TPA targeting ACS affected acetate metabolism, causing significantly decreased acetate consumption (at both 12 and 24 h of cultivation) compared to those of the two control strains (Fig 3D), simultaneously, the growth of *E. coli* was also impaired with TPA (Fig 3E). However, no obvious transcriptional difference of *acs* was observed between the *E. coli* strain containing TPA and the control strain (At2-dCas12a, no crRNA) (Fig 3F), suggesting that the binding of the At2-dCas12a fusion protein to the non-template DNA strand did not affect *acs* transcription. As for RNase II, it has been reported that some mRNAs (such as *ompA* and *udp*) are the direct substrates of this enzyme (32). RNase II can stabilize these mRNAs by removing their poly(A) tails, however, its activity would be impaired with increased acetylation level (31). Thus, we compared the mRNA levels of *ompA* and *udp* with and without the TPA system. The results showed that in vivo mRNA levels of these two genes both decreased, suggesting a changed activity of RNase II in *E. coli* in the presence of TPA (Fig 3G). Similarly, the *E. coli* strain containing the At2-dCas12a-crRNA_*rnb*_ construct also showed impaired growth compared to the control strains (Fig 3H), but no obvious transcriptional difference in *rnb* was detected (Fig 3I).

To further confirm that the TPA editor did not cause substantial nonspecific acetylation, we compared the global protein acetylation levels of the *E. coli* strains containing an active TPA editor (targeting *acs*), an untargeted construct (dCas12a-At2 without crRNA), and a single dCas12a-crRNA construct (control). We detected an obviously elevated global protein acetylation level of the strain with the untargeted construct compared to that containing the dCas12a-crRNA control construct, whereas no obvious change was observed for the active TPA editor (Fig 3J). In addition, we predicted the potential off-targets of the TPA editor targeting *acs* using the Cas-OFFinder tool (http://www.rgenome.net/cas-offinder/) (Supplementary Figure S5A), and then chose two candidates with the least mismatches (6 mismatches) for test. The results showed that the proteins associated with these two off-targets, i.e., NarL and GadE, did not exhibit obviously increased acetylation levels in the presence of TPA (Supplementary Figure S5B), suggesting that the TPA editor can direct the At2 protein to the chosen target for specific acetylation with minimal off-target activity. Collectively, these results demonstrate that TPA can site-specifically increase the acetylation levels of target proteins in bacteria, enabling rapid identification of acetylation events associated with specific phenotypic outcomes.

### TPA-based mini-library screening identified a crucial acetylation-regulated protein RpoD

Sigma factors, as the crucial parts of RNAP holoenzyme, play important roles in promoter recognition and transcription initiation (33). Despite sigma factors have been found to be acetylated (29,34), whether such a modification affects their function in vivo remains minimally explored (17). Here, we focused on all the annoted sigma factor genes (*fecI, rpoS, rpoD, rpoH, rpoE, rpoN*, and *fliA*) in *E. coli* and constructed a mini TPA library to screen functional sigma factors that are associated with phenotypic changes (strain growth) (Fig 4A). Two or three crRNAs were designed for each sigma factor gene in this library. We found that, among all the tested candidates, strains containing the crRNAs targeting *rpoD* (*rpoD*-R2 or *rpoD*-R3) exhibited the weakest growth capability (Fig 4A and 4B); simultaneously, no obvious transcriptional change was detected for *rpoD* in the presence of the TPA construct (Fig 4C). Next, a 6×His tag was inserted into chromosome at the C-terminal of *rpoD* for protein purification (Supplementary Figure S2), yielding the His-tagged RpoD protein for western blot analysis. The results showed that the acetylation level of RpoD in cells containing the TPA editor was much higher than that from the control strain (At2-dCas12a, no crRNA) (Fig 4D), suggesting that TPA did achieve the in vivo specific acetylaiton of RpoD. Together, all these results indicate a causal relationship between the in vivo RpoD acetylation and cell growth.

**Fig 4.**
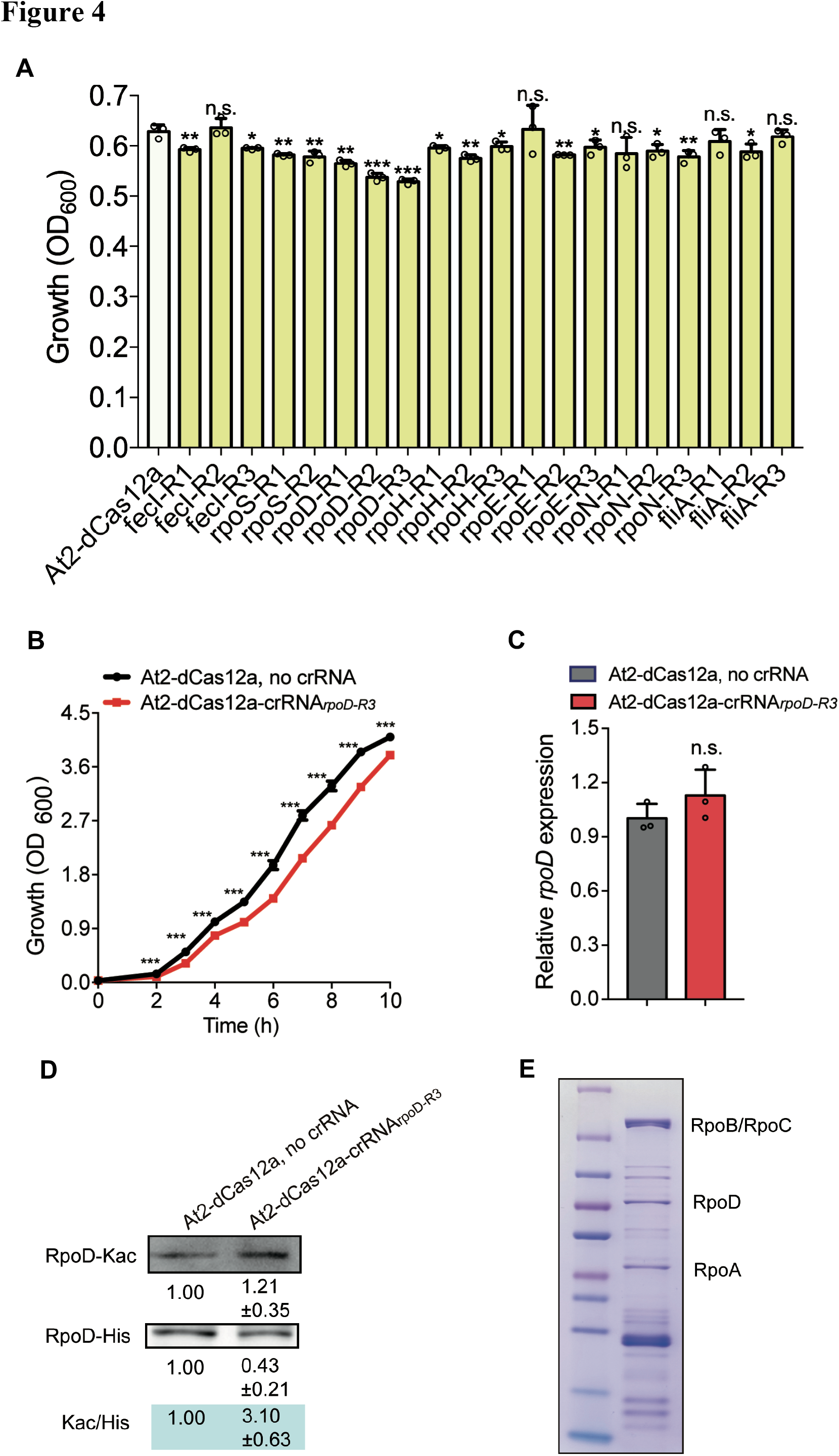
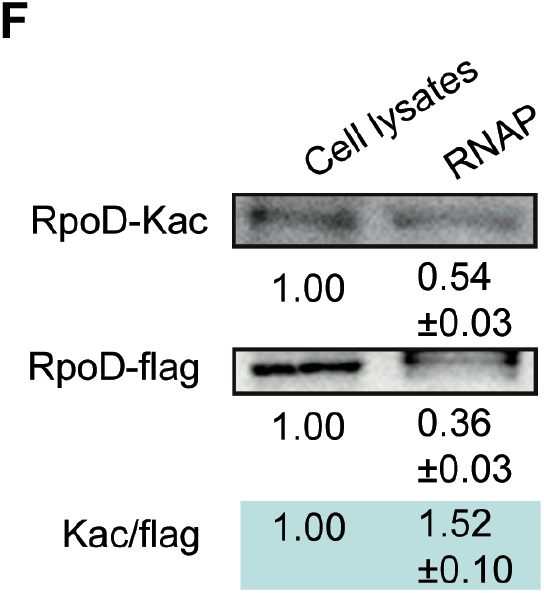
Functional screening of a mini TPA library targeting sigma factors in *E. coli*. (**A**) The growth of *E. coli* cells transferred with the TPA editors targeting different sigma factor genes. The control strain contains an At2-dCas12a fusion without crRNA. Cells were grown at 30°C in LB medium for 3 h. R1, 2, and 3 represent different crRNA-targeting sites of each gene. (**B**) The growth curve of the *E. coli* cells containing the TPA editor targeting RpoD at the R3 site. The control strain contains an At2-dCas12a fusion without crRNA. Cells were grown at 30°C in LB medium for 10h. (**C**) The relative mRNA levels of *rpoD* in the *E. coli* cells containing the TPA editor targeting RpoD at the R3 site and an At2-dCas12a fusion without crRNA. Data are represented as mean ± SD (*n* = 3). Error bars show SDs. Statistical analysis was performed by a two-tailed Student’s *t*-test. ***, *P* < 0.001, **, *P* < 0.01, *, *P* < 0.05, n.s., no significance. (**D**) The in vivo acetylation levels of RpoD with and without the TPA editor. The RpoD proteins in cells containing the TPA editor targeting RpoD and control strain (At2-dCas12a, no crRNA) were analyzed through western bolt with either anti-acetylation (RpoD-Kac) or anti-His (RpoD-His) antibodies. Kac/His, relative ratio of RpoD acetylation in the two strains. The data are presented as mean ± SD from three independent experiments. (**E**) The purified RNAP holoenzyme from the *E. coli* RpoC-6H&RpoD-3F strain (Supplementary Table S1). RpoA, α subunit of RNAP; RpoD, sigma factor of RNAP; RpoB/RpoC, β and β’ subunits of RNAP. (**F**) The acetylation levels of RpoD in the cell lysates and in RNAP from the *E. coli* RpoC-6H&RpoD-3F strain. The RpoD proteins in the cell lysates and in RNAP were analyzed through western bolt with either anti-acetylation (RpoD-Kac) or anti-flag (RpoD-flag) antibodies. Kac/flag, relative ratio of RpoD acetylation in the cell lysates vs. RNAP-bound RpoD. The data are presented as mean ± SD from three independent experiments.

A following question is how acetylation regulates the function of RpoD? It has been known that RpoD, as a sigma factor, can bind to the core enzyme of RNA polymerase (RNAP) and enable the recognition of promoters in *E. coli* (35). Here, we compared the acetylation statuses of RpoD in the cell lysates and in RNAP (Fig 4E). The results showed that the Kac/flag ratio of the RpoD in RNAP was higher (1.52-fold) than that in the cell lysates (Fig 4F), suggesting that the RpoD in RNAP was acetylated to a higher level than in cell lysates. This finding indicates that the increased acetylation level of RpoD may affect its binding to RNAP.

### Targeted protein acetylation in gram-positive bacteria

Following validation of the TPA editor in *E. coli*, we further tested this system in gram-positive bacteria. *C. ljungdahlii*, a representative gram-positive, anaerobic bacterium, was chosen for the assessment.

We constructed a plasmid containing the At2-dCas12a fusion protein, the *adhE1* gene, which is known to be acetylated and responsible for ethanol synthesis in *C. ljungdahlii* (25) (Supplementary Figure S6A), and a designed crRNA targeting the non-template strand of *adhE1* (Fig 5A). A 6×His tag sequence was inserted at the C-terminus of *adhE1* on the plasmid for protein purification. This plasmid was transferred into *C. ljungdahlii* (Δ*dat1*) for the targeted AdhE1 acetylation. The results showed that the expression of At2-dCas12a-crRNA increased the acetylation level of AdhE1 to 1.54-fold of that without crRNA (Fig 5A), suggesting that TPA achieved higher AdhE1 acetylation in *C. ljungdahlii*. Interestingly, the yielding strain showed quite different product synthesis compared to the control strain (without crRNA), despite the fact that their growth was almost identical (Fig 5B and Supplementary Figure S6B). Considering the similar transcriptional levels of *adhE1* in the TPA-contained and control strain (Supplementary Figure S6B), these data suggest a causal relationship between the increased AdhE1 acetylation and changed product synthesis of *C. ljungdahlii*, indicating the utility of TPA in gram-positive bacteria.

**Fig 5.**
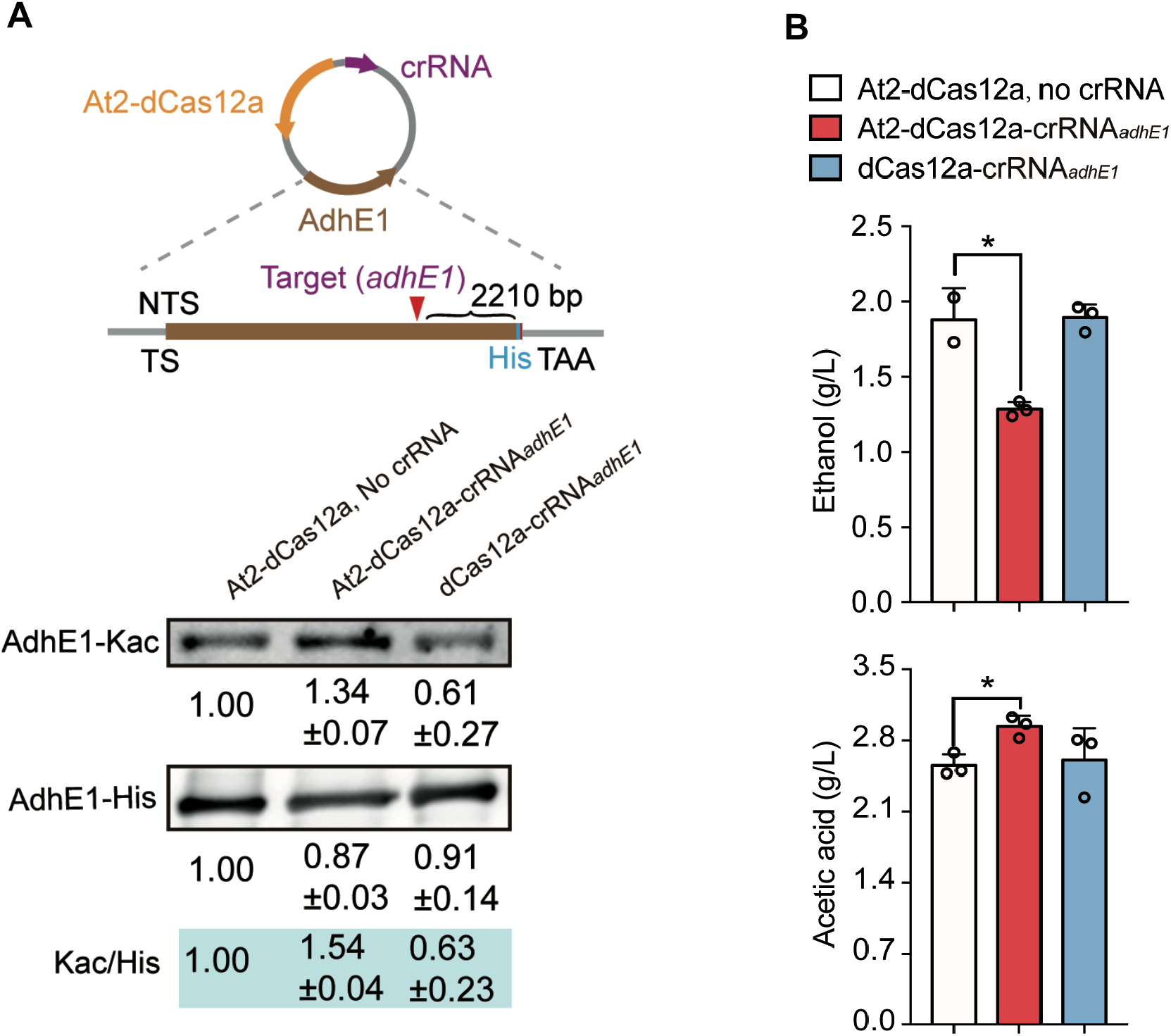
Validation of the TPA editor in *C. ljungdahlii*. (**A**) The plasmid carrying the TPA editor and target gene (*adhE1* from *C. ljungdahlii*) with a 6×His tag at its C-terminus. The plasmid was transferred into *C. ljungdahlii* for in vivo specific acetylation and the His-tagged AdhE1 was then purified for western blot analysis. The acetylation levels of AdhE1 with and without TPA editor. The two controls were the At2-dCas12a fusion (without crRNA) and the single dCas12a (with crRNA). The data are presented as mean ± SD from three independent experiments. (**B**) The product formation of *C. ljungdahlii* cells transferred with the TPA editor, At2-dCas12a fusion (no crRNA), and dCas12a (with crRNA) in gas fermentation. Data are represented as mean ± SD (*n* = 3). Error bars show SDs. Statistical analysis was performed by a two-tailed Student’s *t*-test. *, *P* < 0.05 versus the control (At2-dcas12a, no crRNA).

### Construction and validation of a targeted protein propionylation system in bacteria

After testing and verifying the efficacy of TPA, we examined whether its application could be expanded to other types of acylation modifications, such as targeted protein propionylation (TPP) (Fig 6A). To this end, the At2 enzyme in TPA was replaced by YfiQ, a reported propionyltransferase from *E. coli* (36), yielding a YfiQ-dCas12a fusion protein (Fig 6B). The plasmids expressing the YfiQ-dCas12a fusion, a crRNA, and the *acs* gene (encoding the target Acs protein) were transferred into *E. coli*. Next, the purified Acs protein was then used for western blot analysis with an anti-propionyllysine antibody. We observed substantial propionylation of Acs upon the expression of the TPP construct, which was much higher than that of the control (Fig 6B), suggesting a specific Acs propionylation in *E. coli*. These results confirm that TPA-like targeting systems can be used for other types of in vivo protein modifications.

**Fig 6.**
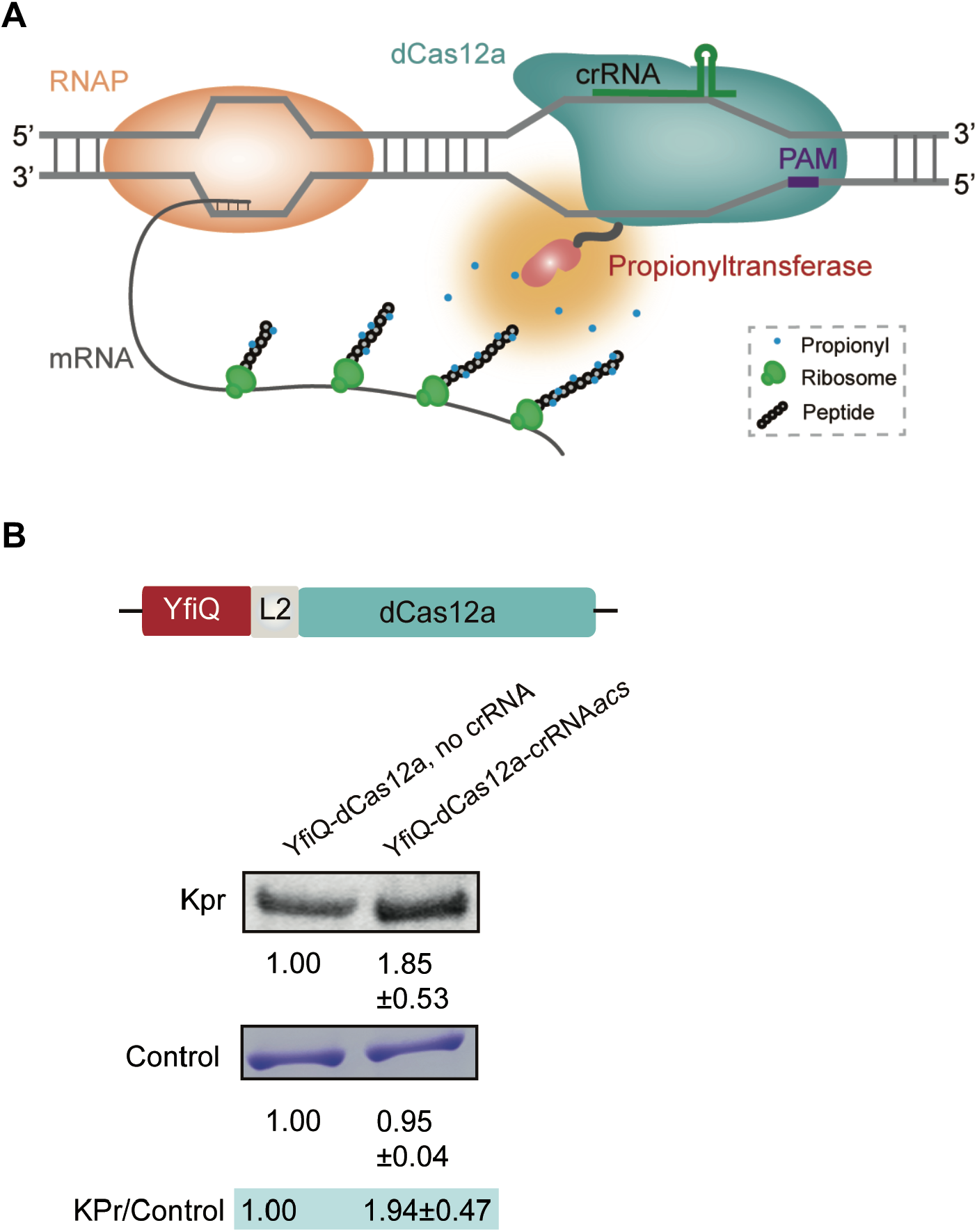
Construction and validation of a targeted protein propionylation (TPP) system in bacteria. (**A**) The proposed mechanism of TPP in bacteria. A fusion protein containing dCas12a and propionyltransferase proteins mediates the crRNA-guided specific propionylation of target proteins based on the synchrony of gene transcription and protein translation in bacteria. (**B**) Plasmids carrying the YfiQ-L2-dCas12a fusion and the *acs* gene were transferred into *E. coli* to test the TPP effect. After targeted propionylation, His-tagged Acs was purified for western blot analysis. The control was the At2-dCas12a fusion (without crRNA). Kpr, lysine (K) propionylation. The data are presented as mean ± SD from three independent experiments.

## DISCUSSION

Tools for the in vivo targeted modification of proteins are currently missing. Their development will enable rapid elucidation of causal relationships between individual protein modification events and phenotypic outcomes. This can facilitate our understanding of the specific functions of protein modifications, and therefore, is crucial for functional proteomics research. In the present study, we developed a system for targeted protein acetylation in bacteria by combining acetyltransferases that are responsible for protein lysine acetylation and the DNA-targeting capability of the CRISPR-Cas12a complex. Our findings demonstrate the specificity and low off-targeting activity of TPA in operation, as well as its potential universality in bacteria.

As to the in vivo targeted modification methods, the earliest reports were about zinc-finger nucleases or transcription activator-like effector based specific methylation or demethylation of DNA and eukaryotic histones (37-39). Furthermore, the target protein deglycosylation and glycosylation by a nanobody-fused N-acetylglucosamine eraser and transferase, respectively, have also been achieved in mammalian cells (40,41). Compared with these editors, CRISPR/Cas-based targeted modifications to nucleic acids or proteins have multiple key advantages, especially in terms of efficiency and convenience. The guide RNA (20□24 bp) is the only component that needs to be changed when using Cas-directed editors for modifications to different targets. This allows the system to easily adapt to various bacterial hosts. To date, Cas9/Cas13-directed DNA/RNA methylation/demethylation and Cas9-directed histone methylation/demethylation have been achieved in bacteria and mammalian cells (42-46). With regard to targeted protein acylation, the only advance has been the Cas9-directed acetylation of histones in eukaryotic cells, increasing the acetylation levels of histones and subsequently altering the expression of histone-associated genes (46). However, due to the asynchrony of gene transcription and protein translation in eukaryotes, the TPA-like in vivo protein modifications to target any proteins of interest have not been achieved and needs to be further studied.

As an efficient targeted protein acetylation tool, the real value of the TPA editor resides in its capacity to rapidly reveal causal relationships between individual protein acetylation events and important phenotypes in bacteria, although some lysine residues of the target protein may be inaccessible to TPA. Therefore, TPA exhibits advantages over the existing methods in the functional examination of specific protein acetylation modifications. As to the potential influence of TPA on gene transcription, we have proved that designing the DNA-binding site of the TPA construct at the 3’ end of the non-template DNA strand of the target gene could avoid this possibility to greatest extent (Fig 3F and 3I). Moreover, it has been found that the dCas12a binding to the non-template strands of target genes had no or minimal transcriptional repression (47,48). Therefore, it can be concluded that the TPA-based identification of causal relationships between individual protein acetylation modifications and phenotypic outcomes is credible.

We found that the acetylation sites of the target protein in the presence of TPA were not entirely faithful to the case without TPA, although such a modification was not significant (Supplementary Table S3). It has been known that protein lysine acetylation is a dynamic and reversible process in cells (1). Thus, this phenomenon should be attributed to the interference of TPA on the natural lysine acetylation process of the target protein. In addition, it should be pointed out that a major limitation of the TPA system is that it cannot target an individual lysine residue; thus, this tool is unable to directly link phenotypic changes to a specific lysine acetylation site of the target protein.

Normally, there are hundreds or even thousands of acetylated proteins generated in the proteomics data of a single bacterial species; but the existing technology can not support an efficient screening to identify the potential crucial protein acetylation events. Through a TPA pool targeting abundant proteins, combined with high-throughput sequencing, it will enable large-scale identification of protein acetylation events associated with important phenotypes in bacteria. The rapid functional screening of the mini TPA library targeting all the sigma factors of *E. coli* in this work also supports this possibility. Furthermore, we have demonstrated targeted acetylation and propionylation in bacteria; in light of these findings, we anticipate that TPA-like editors can be adapted to more targeted protein modifications by employing other modification enzymes (e.g., kinase, methyltransferase, and glycosyltransferase). In addition, although TPA appears to only work in bacteria currently, a further improvement of this system with other Cas proteins would lead to a widespread utility of this tool. For example, acetyltransferases may be guided to mRNA by using a catalytically inactivated Cas13 (dCas13) that binds to ssRNA, which might achieve the use of TPA in eukaryotes. Overall, this study represents a crucial step that guides the development of tools for in vivo targeted protein modification in cells.

## DATA AVAILABILITY

All data supporting the findings of this study are available in the article and its supplementary figures and tables or can be obtained from the corresponding authors upon reasonable request. The mass spectrometry data have been deposited to the ProteomeXchange Consortium (http://proteomecentral.proteomexchange.org) via the iProX partner repository with the dataset identifier PXD031674.

## ACKNOWLEDGEMENTS

We thank Prof. Yong Wang and Prof. Sheng Yang for their kind and generous help. *Author contributions:* Y. L. constructed the TPA system, analyzed the phenotypic changes of mutants, and performed biochemical analyses. N. Z. constructed plasmids, discussed results and offered advices. W. J. and Y. G. supervised and directed the study. Y. L., W. J., and Y. G. wrote the manuscript.

