## Supplementary material for "Programmable acetylation modification of bacterial proteins by a Cas12a-guided acetyltransferase": Supplementary information.pdf

### Supplementary Figures

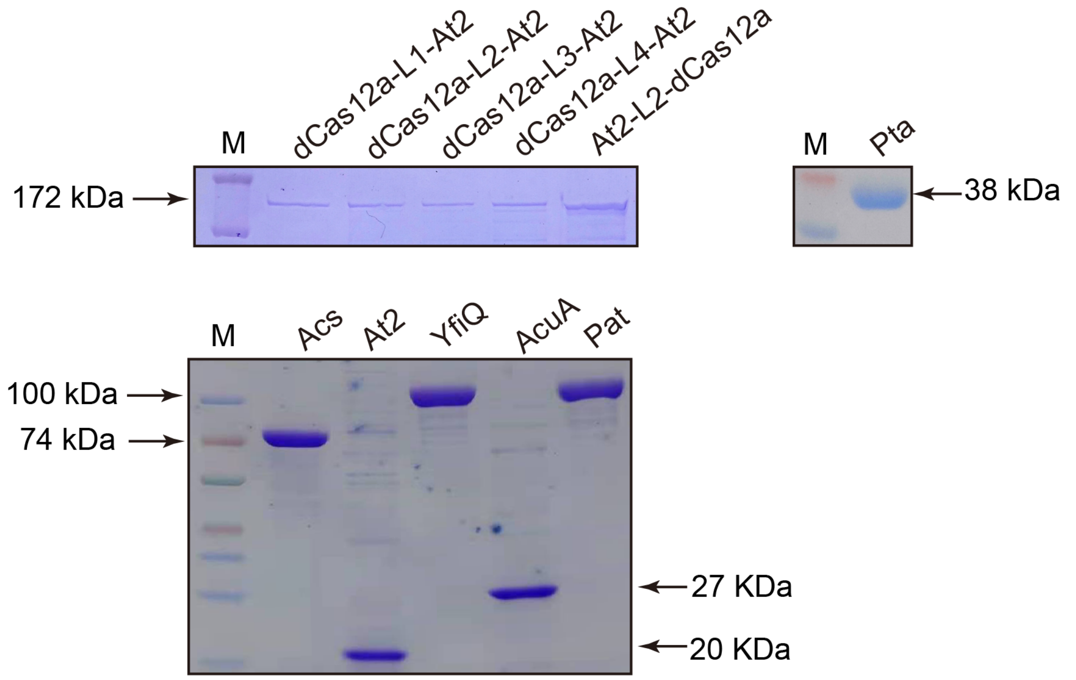


**Supplementary Figure S1 Purification of proteins was determined by SDS-PAGE.** The molecular weight was dCas12a-L1-At2 (172 kDa), dCas12a-L2-At2 (172 kDa), dCas12a-L3-At2 (172 kDa), dCas12a-L4-At2 (172 kDa), At2-L2-dCas12a (172 kDa), Pta (38 kDa), Acs (74 kDa), At2 (20kDa), YfiQ (100 kDa), AcuA (27 kDa), and Pat (100 kDa)**.**

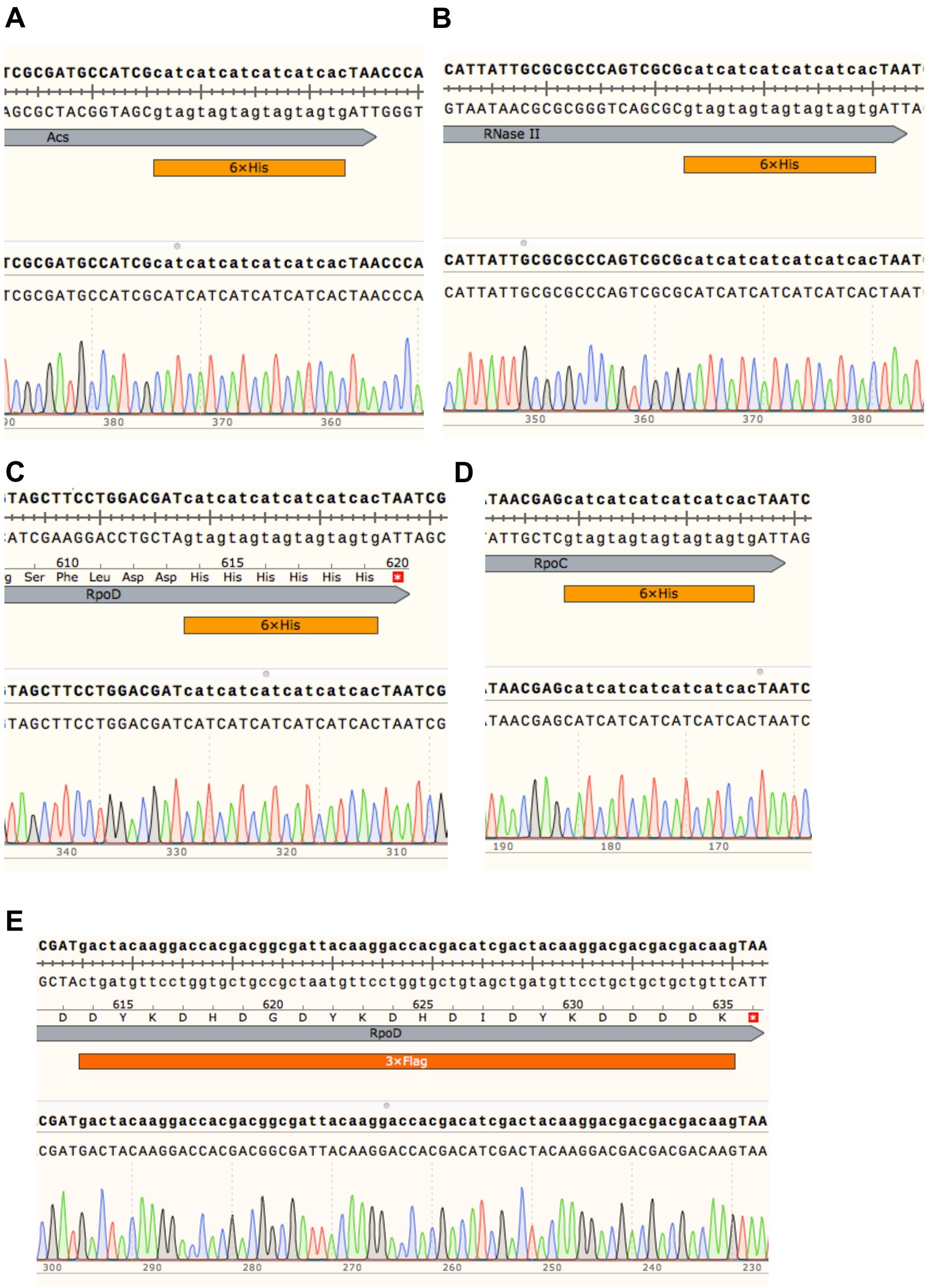


**Supplementary Figure S2 (A-D) Confirmation of the insertion of 6×His tag into *acs*, *rnb, rpoD, and rpoC* by sequencing. (E) Confirmation of the insertion of 3×Flag tag into *rpoD* by sequencing.**

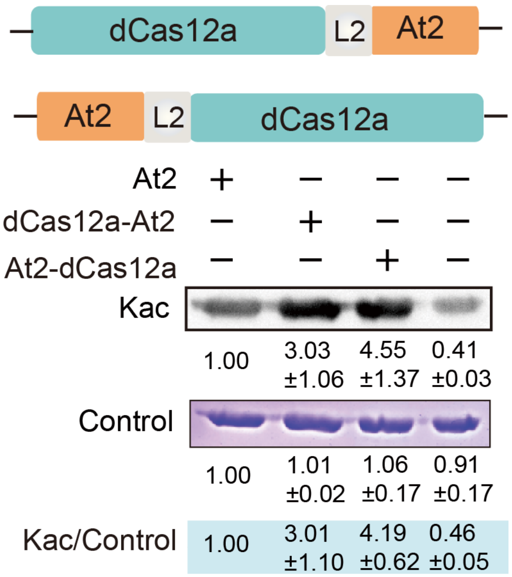


**Supplementary Figure S3 Influence of the relative location of At2 and dCas12a in the fusion protein on At2 activity.**

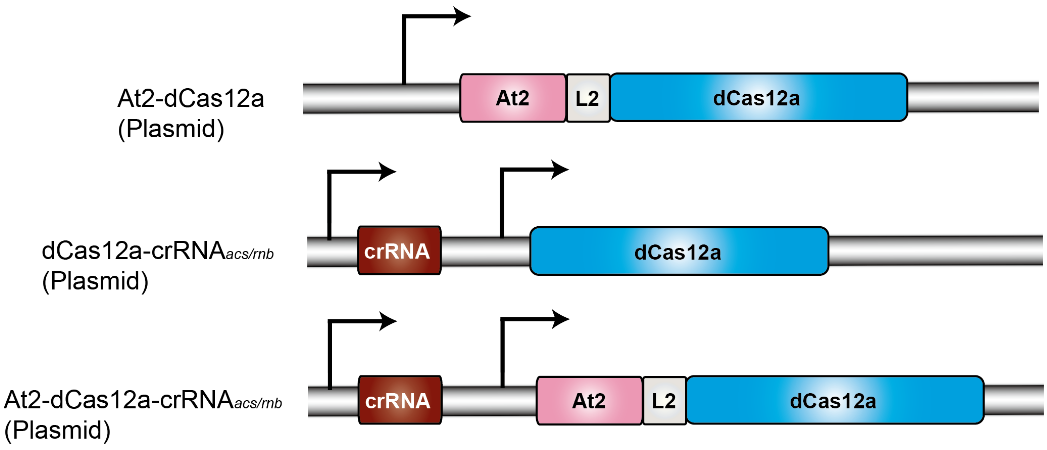


**Supplementary Figure S4 Plasmids designed to explore the phenotypic changes caused by altering of acetylation levels on Acs and RNase II.**

**A**

Summary

| Target Sequence | Bulge Type | Bulge Size | Mismatch | Number of Found Targets |
|---|---|---|---|---|
| TTNCGCAGAATACGGCGCATAATTTTG | X | 0 | 0 | 1 |
| TTNCGCAGAATACGGCGCATAATTTTG | X | 0 | 6 | 2 |
| TTNCGCAGAATACGGCGCATAATTTTG | X | 0 | 7 | 4 |
| TTNCGCAGAATACGGCGCATAATTTTG | X | 0 | 8 | 28 |
| TTNCGCAGAATACGGCGCATAATTTTG | X | 0 | 9 | 115 |

| crRNA | DNA | Position | Direction | Mismatches |
|---|---|---|---|---|
| TTNCGCAGAATACGGCGCATAATTTTG | TTGCGCAGAATACGGCGCATAATTTTG | 356104 | - | 0 |
| TTNCGCAGAATACGGCGCATAATTTTG | TTTttCAGAATcaGttGCATAATTTTG (*gadE*) | 982898 | + | 6 |
| TTNCGCAGAATACGGCGCATAATTTTG | TTGCGaAGAgTACaGCGCcaAATTTgG (*narL*) | 3365437 | + | 6 |
| TTNCGCAGAATACGGCGCATAATTTTG | TTAaaCAGAAaAtGGgGatTAATTTTG | 2141858 | + | 7 |
| TTNCGCAGAATACGGCGCATAATTTTG | TTGacgAaAATcaGGCGCAaAATTTTG | 3256030 | - | 7 |
| TTNCGCAGAATACGGCGCATAATTTTG | TTTCGCcGgATgCatCGCAaAATTgTG | 3818237 | + | 7 |
| TTNCGCAGAATACGGCGCATAATTTTG | TTTCGCAGcATcCGGCaCtTAtTgTcG | 3835752 | - | 7 |
| TTNCGCAGAATACGGCGCATAATTTTG | TTAtGCAGcAcACGcgcCATccTTTTG | 79365 | + | 8 |
| TTNCGCAGAATACGGCGCATAATTTTG | TTCacgAagATAtGGCGaAgAATTTTG | 504272 | + | 8 |
| TTNCGCAGAATACGGCGCATAATTTTG | TTCCaCAGcATAtGGttggTAATgTTG | 254273 | - | 8 |

**B**

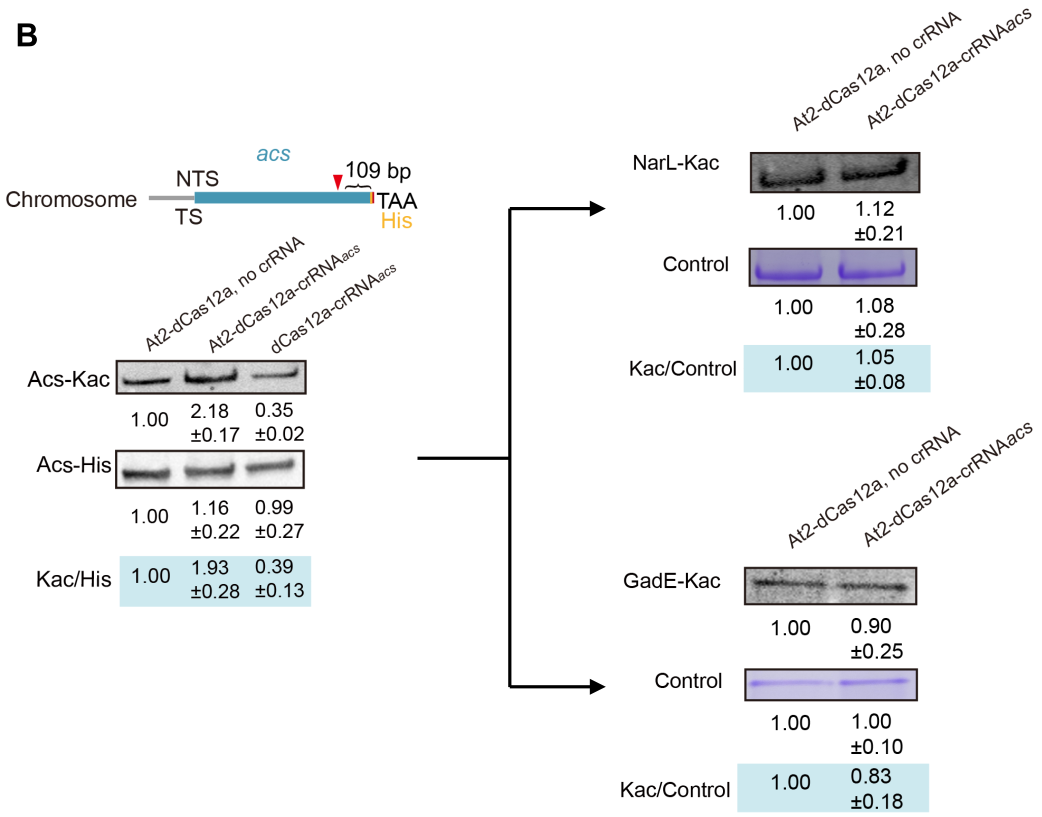

**Supplementary Figure S5 Evaluate the off-target activity of TPA.** (**A**) Prediction of potential off-targets of the TPA editor (targeting *acs*) using the Cas-OFFinder tool. The two off-targets (highlighted with red) that have six mismatched bases against the dCas12a-At2-crRNA*acs*-trageted DNA sequence were subjected to the following analysis. (**B**) Left: TPA binding to *acs* at the chromosome and acetylation levels of Acs with and without TPA editor. Right: TPA binding to the potential off-targets (*narL* and *gadE* expressed on plasmid) and acetylation levels of NarL and GadE with and without TPA editor. Cells were grown at 37°C in LB medium. The data are presented as mean ± SD ($n$ = 2).

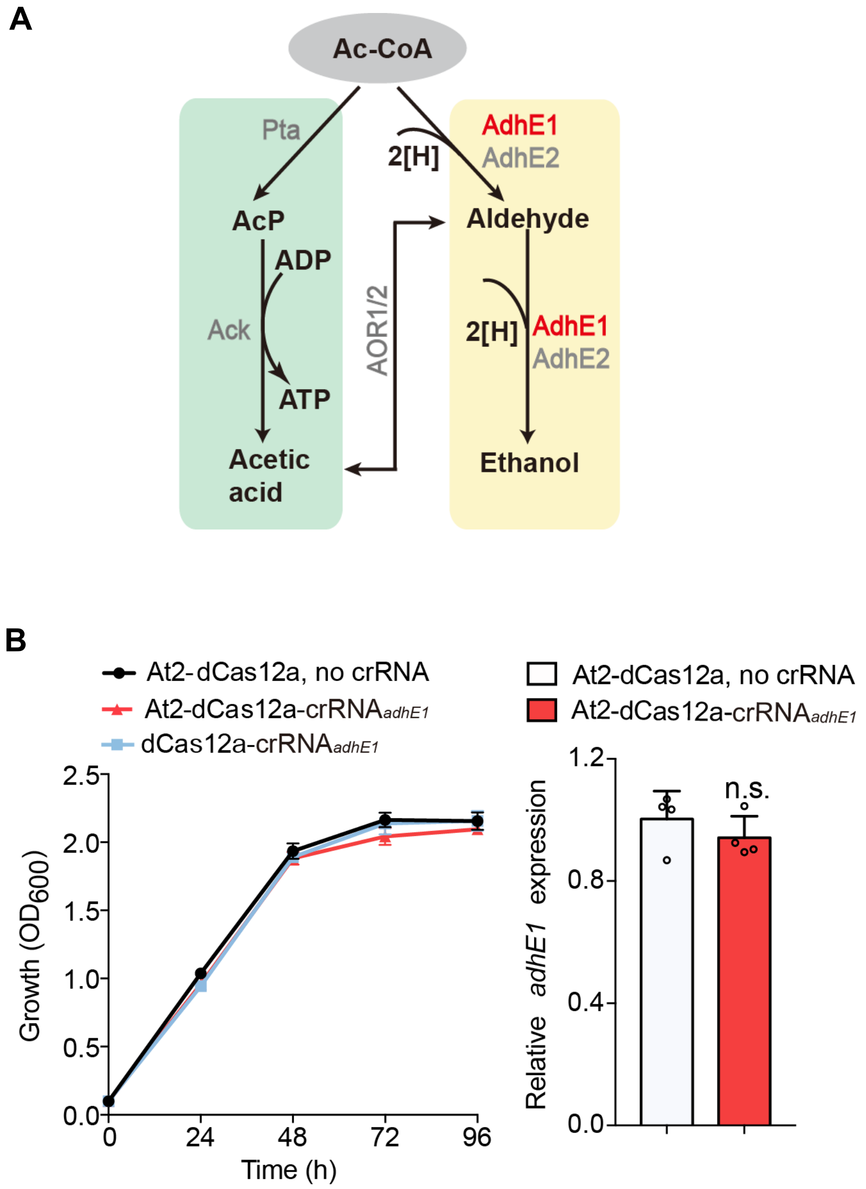


**Supplementary Figure S6 Targeted AdhE1 acetylation in *Clostridium ljungdahlii*.** (**A**) Schematic diagram of acetic acid and ethanol production as well as their conversion in *C. ljungdahlii*. (**B**) Comparison of the growth of *C. ljungdahlii* strains transferred with TPA editor, At2-dCas12a fusion (no crRNA), or dCas12a (with crRNA) in gas fermentation. Comparison of the *adhE1* transcript levels of *C. ljungdahlii* strains transferred with TPA editor and At2-dCas12a fusion (no crRNA) in gas fermentation. Data are represented as mean ± SD ($n$ = 3). Error bars show SDs. Statistical analysis was performed by a two-tailed Student's $t$-test. n.s., no significant changes versus the control (At2-dcas12a, no crRNA).

### Supplementary Tables

#### Supplementary Table S1: Strains and plasmids used in this study

| **Strains or plasmids** | **Description of genotype** | **Sources or references** |
|---|---|---|
| **Bacterial strains** | | |
| ***C. ljungdahlii*** | | |
| WT | DSM 13528,wild-type strain | DSMZ |
| ΔDat1 | DSM 13528, Δ*dat1* (CLJU_c01320) | This lab |
| ***E. coli*** | | |
| BL21 (DE3) | Strain used for protein overexpression | Novagen |
| DH5α | General cloning host strain | Invitrogen |
| BW25113 | Strain used for phenotype research | Novagen |
| Acs-6H | BW25113, inserted a 6×His tag at C-termianl of *acs* | This study |
| Rnb-6H | BW25113, inserted a 6×His tag at C-termianl of *rnb* | This study |
| RpoD-6H | BW25113, inserted a 6×His tag at C-termianl of *rpoD* | This study |
| RpoC-6H&RpoD-3F | BW25113, inserted a 6×His tag at C-termianl of *rpoC* and a 3×Flag tag at C-termianl of *rpoD* | This study |
| **Plasmids** | | |
| pACYCDuet-1 | p15A ori, $Cm^R$ | Gift from Prof. Yong Wang |
| pMTL83151 | pCB102 ori, $Cm^R$ | This lab |
| pET-28a | pBR322 ori, $Kan^R$ | This lab |
| pMTL83151-Dc-at2-adhE1 (6H) | dCas12a-L2-at2 & AdhE1-6×His expression vector, derived from pMTL83151 | This study |

| | | |
|---|---|---|
| pMTL83151-Dc-at2-crRNA$_{adhE1}$-adhE1 (6H) | dCas12a-L2-at2- crRNA$_{adhE1}$ & AdhE1-6×His expression vector, derived from pMTL83151 | This study |
| pMTL83151-Dc-crRNA$_{adhE1}$-adhE1 (6H) | dCas12a- crRNA$_{adhE1}$ & AdhE1-6×His expression vector, derived from pMTL83151 | This study |
| pET-28a-Ndc-L1-At2 | pET-28a carrying the *dCas12a-l1-at2* gene cluster, derived from pET-28a | This study |
| pET-28a-Ndc -L2-At2 | pET-28a carrying the *dCas12a-l2-at2* gene cluster, derived from pET-28a | This study |
| pET-28a-Ndc-L3-At2 | pET-28a carrying the *dCas12a-l3-at2* gene cluster, derived from pET-28a | This study |
| pET-28a-Ndc-L4-At2 | pET-28a carrying the *dCas12a-l4-at2* gene cluster, derived from pET-28a | This study |
| pET-28a-Nat2-L2-Dc | pET-28a carrying the *the at2-l2-dCas12a gene cluster* cluster, derived from pET-28a | This study |
| pNat2-L2-Dc-NCr | pACYC carrying the *the at2-l2-dCas12a gene cluster* cluster, derived from pACYCDuet-1 | This study |
| pDc-acs-Ts | pACYC carrying the *dCas12a* gene and crRNA named *acs*-Ts, derived from pACYCDuet-1 | This study |
| pNat2-L2-Dc-acs-Ts | pACYC carrying the the at2-l2-dCas12a gene cluster cluster and crRNA named *acs*-Ts, derived from pACYCDuet-1 | This study |
| pDc-rnb-Ts | pACYC carrying the *dCas12a* gene and crRNA named *rnb*-Ts, derived from pACYCDuet-1 | This study |
| pNat2-L2-Dc-rnb-Ts | pACYC carrying the the at2-l2-dCas12a gene cluster cluster and crRNA named *rnb*-Ts, derived from pACYCDuet-1 | This study |

| | | |
|---|---|---|
| pET-28a-pta | pET-28a carrying the *pta* gene, derived from pET-28a | This study |
| pET-28a-acs | pET-28a carrying the *acs* gene, derived from pET-28a | This study |
| pET-28a-lrp | pET-28a carrying the *lrp* gene, derived from pET-28a | This study |
| pET-28a-rpoD | pET-28a carrying the *rpoD* gene, derived from pET-28a | This study |
| pNat2-L2-Dc-acs-NTs | pACYC carrying the the at2-l2-dCas12a gene cluster cluster and crRNA named *acs*-NTs, derived from pACYCDuet-1 | This study |
| pDc-acs-NTs | pACYC carrying the *dCas12a* gene and crRNA named *acs*-NTs, derived from pACYCDuet-1 | This study |
| pNat2-L2-Dc-lrp-Ts1 | pACYC carrying the the at2-l2-dCas12a gene cluster cluster and crRNA named *lrp*-Ts1, derived from pACYCDuet-1 | This study |
| pNat2-L2-Dc-lrp-Ts2 | pACYC carrying the the at2-l2-dCas12a gene cluster cluster and crRNA named *lrp*-Ts2, derived from pACYCDuet-1 | This study |
| pNat2-L2-Dc-rnb-NTs | pACYC carrying the the at2-l2-dCas12a gene cluster cluster and crRNA named *rnb*-NTs, derived from pACYCDuet-1 | This study |
| pDc-rnb-NTs | pACYC carrying the *dCas12a* gene and crRNA named *rnb*-NTs, derived from pACYCDuet-1 | This study |
| pNat2-L2-Dc-fecI-R1 | pACYC carrying the the at2-l2-dCas12a gene cluster cluster and crRNA named *fecI*-R1, derived from pACYCDuet-1 | This study |
| pNat2-L2-Dc-fecI-R2 | pACYC carrying the the at2-l2-dCas12a gene cluster cluster and crRNA named *fecI*-R2, derived from pACYCDuet-1 | This study |

| | | |
|---|---|---|
| pNat2-L2-Dc-fecI-R3 | pACYC carrying the the at2-l2-dCas12a gene cluster cluster and crRNA named *fecI*-R3, derived from pACYCDuet-1 | This study |
| pNat2-L2-Dc-rpoS-R1 | pACYC carrying the the at2-l2-dCas12a gene cluster cluster and crRNA named *rpoS*-R1, derived from pACYCDuet-1 | This study |
| pNat2-L2-Dc-rpoS-R2 | pACYC carrying the the at2-l2-dCas12a gene cluster cluster and crRNA named *rpoS*-R2, derived from pACYCDuet-1 | This study |
| pNat2-L2-Dc-rpoD-R1 | pACYC carrying the the at2-l2-dCas12a gene cluster cluster and crRNA named *rpoD*-R1, derived from pACYCDuet-1 | This study |
| pNat2-L2-Dc-rpoD-R2 | pACYC carrying the the at2-l2-dCas12a gene cluster cluster and crRNA named *rpoD*-R2, derived from pACYCDuet-1 | This study |
| pNat2-L2-Dc-rpoD-R3 | pACYC carrying the the at2-l2-dCas12a gene cluster cluster and crRNA named *rpoD*-R3, derived from pACYCDuet-1 | This study |
| pNat2-L2-Dc-rpoH-R1 | pACYC carrying the the at2-l2-dCas12a gene cluster cluster and crRNA named *rpoH*-R1, derived from pACYCDuet-1 | This study |
| pNat2-L2-Dc-rpoH-R2 | pACYC carrying the the at2-l2-dCas12a gene cluster cluster and crRNA named *rpoH*-R2, derived from pACYCDuet-1 | This study |
| pNat2-L2-Dc-rpoH-R3 | pACYC carrying the the at2-l2-dCas12a gene cluster cluster and crRNA named *rpoH*-R3, derived from pACYCDuet-1 | This study |
| pNat2-L2-Dc-rpoE-R1 | pACYC carrying the the at2-l2-dCas12a gene cluster cluster and crRNA named *rpoE*-R1, derived from pACYCDuet-1 | This study |

| | | |
|---|---|---|
| pNat2-L2-Dc-rpoE-R2 | pACYC carrying the at2-l2-dCas12a gene cluster and crRNA named *rpoE*-R2, derived from pACYCDuet-1 | This study |
| pNat2-L2-Dc-rpoE-R3 | pACYC carrying the at2-l2-dCas12a gene cluster and crRNA named *rpoE*-R3, derived from pACYCDuet-1 | This study |
| pNat2-L2-Dc-rpoN-R1 | pACYC carrying the at2-l2-dCas12a gene cluster and crRNA named *rpoN*-R1, derived from pACYCDuet-1 | This study |
| pNat2-L2-Dc-rpoN-R2 | pACYC carrying the at2-l2-dCas12a gene cluster and crRNA named *rpoN*-R2, derived from pACYCDuet-1 | This study |
| pNat2-L2-Dc-rpoN-R3 | pACYC carrying the at2-l2-dCas12a gene cluster and crRNA named *rpoN*-R3, derived from pACYCDuet-1 | This study |
| pNat2-L2-Dc-filA-R1 | pACYC carrying the at2-l2-dCas12a gene cluster and crRNA named *filA*-R1, derived from pACYCDuet-1 | This study |
| pNat2-L2-Dc-filA-R2 | pACYC carrying the at2-l2-dCas12a gene cluster and crRNA named *filA*-R2, derived from pACYCDuet-1 | This study |
| pNat2-L2-Dc-filA-R3 | pACYC carrying the at2-l2-dCas12a gene cluster and crRNA named *filA*-R3, derived from pACYCDuet-1 | This study |
| pTargetF-N20-acs | pMB1 ori, spec$^R$, gRNA of gene *acs* | This study |
| pEcCas | pSC101 ori, Kan$^R$ | Gift from Prof. Sheng Yang |
| pTargetF-N20-rnb | pMB1 ori, spec$^R$, gRNA of gene *rnb* | This study |
| pTargetF-N20-rpoD | pMB1 ori, spec$^R$, gRNA of gene *rpoD* | This study |
| pTargetF-N20-rpoC | pMB1 ori, spec$^R$, gRNA of gene *rpoD* | This study |

| | | |
|---|---|---|
| pNYfiQ-L2-Dc-acs-NTs | pACYC carrying YfiQ-l2-*dCas12a* gene and crRNA named *acs*-NTs, derived from pACYCDuet-1 | This study |

**Supplementary Table S2: Primers used in this study.**

| Primer name | Sequence (5′-3′) |
|---|---|
| $crRNA_{adhE1}$-F | CAATTTCTACTGTTGTAGATTACCATTTCTAGTTTTTAAAGCT AATAAAAATAAGAAG |
| $crRNA_{adhE1}$-R | GATAAAAATAAGAAGCCTGCAAATGCAGGCTTCTTATTTTTA TTAGCTTTAAAAACTA |
| 83151-crRNA-F | CCCGTATCAAAATTTAGGAGGTTAGGATCCAATTTCTACTGT TGTAGAT |
| 83151-crRNA-R | ATGTCTGCAGGCCTCGACCCCGGGCCATGGATAAAAATAAG AAGCCTGCAAATGCAGGC |
| Pet-dCas-F | GCAGCGGCCTGGTGCCGCGCGGCAGCCATATGATGAGTATA TATCAAGAATTTGTT |
| Pet-dCas-L1-R | TCTTATCATGGAGCCACCGCCATTATTTCTATTTTGTACAAAT TC |
| Pet-L1-at2-F | AGAAATAATGGCGGTGGCTCCATGATAAGAAGAGGAAATAT AAAAG |
| Pet-at2-R | TAGTTAAGTATAAGAAGGAGATATACATATGATGATAAGAAG AGGAAATATAAAAG |
| Pet-dCas-L2-R | TTTGGCAGCGGCTTCACCGGAGCCATTATTTCTATTTTGTAC AAATTC |
| Pet-L2-at2-F | GGCTCCGGTGAAGCCGCTGCCAAAATGATAAGAAGAGGAA ATATAAAAG |
| Pet-dCas-L3-R | TCATTTTGGCAGCGGCTTCTTTGGCAGCGGCTTCACCGGAG CCATTATTTCTATTTTGTACAAATTCAAAATATTC |
| Pet-L3-at2-F | ATAATGGCTCCGGTGAAGCCGCTGCCAAAGAAGCCGCTGCC AAAATGATAAGAAGAGGAAATATAAAAGATC |
| Pet-dCas-L4-R | CATTTTGGCAGCGGCTTCTTTGGCAGCGGCTTCTTTGGCAG CGGCTTCACCGGAGCCATTATTTCTATTTTGTACAAATTCAA AA |

| | |
|---|---|
| Pet-L4-at2-F | AATGGCTCCGGTGAAGCCGCTGCCAAAGAAGCCGCTGCCAAAGAAGCCGCTGCCAAAATGATAAGAAGAGGAAATATAAAAG |
| Pet-at2-L2-F | CCTGGTGCCGCGCGGCAGCCATATGATGATAAGAAGAGGAAATATAAAAG |
| Pet-at2-L2-R | ACTCATTTTGGCAGCGGCTTCACCGGAGCCTTTTTAATTTCATACATTAGATAATGCG |
| Pet-L2-dCas-F | AAAAGGCTCCGGTGAAGCCGCTGCCAAAATGAGTATATATCAAGAATTTGTTAATAAAT |
| Pet-L2-dCas-R | ATCTCAGTGGTGGTGGTGGTGGTGCTCGAGTCAATTATTTCTATTTTGTACAAATTCAA |
| At2-F | AGTATAAGAAGGAGATATACATATGATGATAAGAAGAGGAAATATAAAAGATC |
| At2-L2-R | GCTCATTTTGGCAGCGGCTTCACCGGAGCCTTTTTAATTTCATACATTAGATAATGCG |
| dCas12a-L2-F | AATTAAAAAGGCTCCGGTGAAGCCGCTGCCAAAATGAGCATCTATCAGGAGTTCG |
| dCas12a-R | CAGCGGTTTCTTTACCAGACTCGAGTTAGTTGTTACGGTTTTGAACGAATTC |
| crRNA-F | AACTTTAATAAGGAGATATACCATGGAATTTCTACTGTTGTAGAT |
| crRNA-R | GCGCGCCGAGCTCGAATTCGGATCCATAAAAATAAGAAGCCTGCAAA |
| crRNAacs-Ts-F | GAATTTCTACTGTTGTAGATGCCCGCTGGCGACGCCAGACGTGCATAAAAATAAGAAG |
| crRNAacs-Ts-R | CATAAAAATAAGAAGCCTGCAAATGCAGGCTTCTTATTTTTATGCACGTCTGGCGTCG |
| crRNAacs-NTs-F | GAATTTCTACTGTTGTAGATCGCAGAATACGGCGCATAATTTTGATAAAAATAAGAAG |
| crRNAacs-NTs-R | CATAAAAATAAGAAGCCTGCAAATGCAGGCTTCTTATTTTTATCAAAATTATGCGCCG |

| | |
|---|---|
| crRNArnb-NTs-F | GAATTTCTACTGTTGTAGATATCATGTCGCCATATTTACGGATCATAAAAATAAGAAG |
| crRNArnb-NTs-R | CATAAAAATAAGAAGCCTGCAAATGCAGGCTTCTTATTTTTATGATCCGTAAATATGG |
| pET28a-Pta-F | CCTGGTGCCGCGCGGCAGCCATATGATGAAATTGATGGAAAAAATTTGGA |
| pET28a-Pta-R | GCCGGATCTCAGTGGTGGTGGTGGTGGTGCTCGAGTTACTTTTGAGCTTGTGCTTGAAC |
| pET28a-RpoD-F | AGCGGCCTGGTGCCGCGCGGCAGCCATATGATGGAGCAAAACCCGCAGTCACA |
| pET28a-RpoD-R | GCCGGATCTCAGTGGTGGTGGTGGTGGTGCTCGAGTTAATCGTCCAGGAAGCTACGCA |
| acs-Ts-F | GAATTTCTACTGTTGTAGATACATCCTACAAGGAGAACAAAAGCATAAAAATAAGAAGC |
| acs-Ts-R | CATAAAAATAAGAAGCCTGCAAATGCAGGCTTCTTATTTTTATGCTTTTGTTCTCCTTG |
| rnb-Ts-F | GAATTTCTACTGTTGTAGATTGTTTCAGGACAACCCGCTGCTAGATAAAAATAAGAAGC |
| rnb-Ts-R | CATAAAAATAAGAAGCCTGCAAATGCAGGCTTCTTATTTTTATCTAGCAGCGGGTTGTC |
| pET28a-acs-F | GCGGCCTGGTGCCGCGCGGCAGCCATATGATGAGCCAAATTCACAAACACACC |
| pET28a-acs-R | GCCGGATCTCAGTGGTGGTGGTGGTGGTGCTCGAGTTACGATGGCATCGCGATAGCCTG |
| pET28a-At2-F | GCCTGGTGCCGCGCGGCAGCCATATGATGATAAGAAGAGGAAATATAAAAGATC |
| pET28a-At2-R | ATCTCAGTGGTGGTGGTGGTGGTGCTCGAGTTATTTTTTAATTTCATACATTAGATAAT |
| pET28a-AcuA-F | GCGGCCTGGTGCCGCGCGGCAGCCATATGGTGGAACATCATAAAACATACCATTC |
| pET28a-AcuA-R | GATCTCAGTGGTGGTGGTGGTGGTGCTCGAGTTAATACATATAACGATGATAAAAACGG |

| | |
|---|---|
| pET28a-YfiQ-F | ATGGATAAAAAAAAGAATATATCAATGGCTGTTATACAAAGATTGCC |
| pET28a-YfiQ-R | CCGGATCTCAGTGGTGGTGGTGGTGGTGCTCGAGTCATGATTCCTCGCGCTGGGCAAG |
| pET28a-Pat-F | CAGCGGCCTGGTGCCGCGCGGCAGCCATATGATGAGCCAGCAAGGACTGGAAG |
| pET28a-Pat-R | CCGGATCTCAGTGGTGGTGGTGGTGGTGCTCGAGTCACGATTCATCACATTTGGCCAG |
| acs-6H-F | CTAGGTATAATACTAGTCGTCACGCTTAATCACGGGGGTTTTAGAGCTAGAAATAGCA |
| acs-6H-R | TATTTCTAGCTCTAAAACCCCCGTGATTAAGCGTGACGACTAGTATTATACCTAGGACT |
| acs-up-F | GGCGATCACGAACGTTTTGAACAGACCT |
| acs-up-R | TGGTGACGGTTCCTCGCCATGGTTTAGAGTTACATAGGCGTAGATCGCCTGACCTTTAA |
| acs-dw-F | TCTACGCCTATGTAACTCTAAACCATGGCGAGGAACCGTCACCAGAACTGTACGC |
| acs-dw-R | CTAGGTATAATACTAGTCGTCACGCTTAATCACGGGGGTTTTAGAGCTAGAAATAGCA |
| acs-6H-F | CTAGGTATAATACTAGTCGTCACGCTTAATCACGGGGGTTTTAGAGCTAGAAATAGCA |
| acs-6H-R | TATTTCTAGCTCTAAAACCCCCGTGATTAAGCGTGACGACTAGTATTATACCTAGGACT |
| acs-up-F | GGCGATCACGAACGTTTTGAACAGACCT |
| acs-up-R | TGGTGACGGTTCCTCGCCATGGTTTAGAGTTACATAGGCGTAGATCGCCTGACCTTTAA |
| acs-dw-F | TCTACGCCTATGTAACTCTAAACCATGGCGAGGAACCGTCACCAGAACTGTACGC |
| acs-dw-R | CTAGGTATAATACTAGTCGTCACGCTTAATCACGGGGGTTTTAGAGCTAGAAATAGCA |
| rnb-6H-F | AGTCCGCAGCATTATTGCGCGCCGTTTTAGAGCTAGAAATAGCAAGTTAAAATA |

| | |
|---|---|
| rnb-6H-R | CTAAAACGGCGCGCAATAATGCTGCGGACTAGTATTATACCTAGGACTGAGCTAG |
| rnb-up-F | GAACGTGATGTTGGTGACTGGTTATACG |
| rnb-up-R | AGATTAGTGATGATGATGATGATGCGCGACTGGGCGCGCAATAATGCTGCGGGTTTC |
| rnb-dw-F | GCGCGCCCAGTCGCGCATCATCATCATCATCACTAATCTCCTTTCACGGCCCATTCCT |
| rnb-dw-R | TTGTACAACGTTGTGGACTCCCTAACGGT |
| rpoD-6H-F | ACTAGTTGGAGAACTTGTAACCACGGGTTTTAGAGCTAGAAATAGCAAGTTA |
| rpoD-6H-R | TCTAGCTCTAAAACCCGTGGTTACAAGTTCTCCAACTAGTATTATACCTAGGAC |
| rpoD-up-F | GATGAACAAGCCGTGGTCGGAAAA |
| rpoD-up-R | CAGGTTGCGTAGGTAGAAAATTTATATCCGCGACGGTATTCGAATTTATCAACCGCTTT |
| rpoD-dw-F | TTCGAATACCGTCGCGGATATAAATTTTCTACCTACGCAACCTGGTGGATCCGTCAG |
| rpoD-dw-R | TGATCCGGCCTACCGATTAGTGATGATGATGATGATGATCGTCCAGGAAGCTACGCAGC |
| rpoC-6H-F | AATACTAGTGGCGACGCATACGATCCTGGGTTTTAGAGCTAGAAATAGCAAGTTAAAAT |
| rpoC-6H-R | CTAGCTCTAAAACCCAGGATCGTATGCGTCGCCACTAGTATTATACCTAGGACTGAGCT |
| rpoC-up-F | CGTTAACGCGGGTAGCTCCGACTTC |
| rpoC-up-R | TTCACCCGCAGCACGACGGCGCATTCGGTCTTGATGGTACGCGTAACCGGTACCTGCCG |
| rpoC-dw-F | CGCGTACCATCAAGACCGAATGCGCCGTCGTGCTGCGGGTGAAGCTCCGGCTGCACCGC |
| rpoC-dw-R | TTGCGGATTAACGATTAGTGATGATGATGATGATGCTCGTTATCAGAACCGCCCAGACC |
| rpoD-3F-F | ATACTAGTAGCTACGCAGCACTTCAGAAGTTTTAGAGCTAGAAATAGCAAGTTAAAAT |

| | |
|---|---|
| rpoD-3F-R | CTAGCTCTAAAACTTCTGAAGTGCTGCGTAGCTACTAGTATT ATACCTAGGACT |
| rpoD-3F-up-F | GAGCCAATCTCCATGGAAACGCCGATCG |
| rpoD-3F-up-R | TAGTCATCGTCCAGGAAACTTCGAAGGACCTCTGAGCGGCT CGGGTGACGCAGTTTGCG |
| 3F-F | CTCAGAGGTCCTTCGAAGTTTCCTGGACGATGACTACAAGG ACCACGACGGCGA |
| 3F-R | GTGCGGCGTAACGCCTGATCCGGCCTACCGATTACTTGTCGT CGTCGTCCTTGTAGTC |
| rpoD-3F-dw-F | GACGACAAGTAATCGGTAGGCCGGATCAGGCGTTACGCCGC ACCCGGCACTAGG |
| rpoD-3F-dw-R | ACAGTGGGGGAAACAAACGCTCAC |
| fecI-R1-F | GAATTTCTACTGTTGTAGATCGAAAGCAGAAACGCTTCACG TGTATAAAAATAAGAAGC |
| fecI-R1-R | CATAAAAATAAGAAGCCTGCAAATGCAGGCTTCTTATTTTTA TACACGTGAAGCGTTTC |
| fecI-R2-F | GAATTTCTACTGTTGTAGATTGCGCAATCTCGCTGTATGTCA GAATAAAAATAAGAAGC |
| fecI-R2-R | CATAAAAATAAGAAGCCTGCAAATGCAGGCTTCTTATTTTTA TTCTGACATACAGCGAG |
| fecI-R3-F | GAATTTCTACTGTTGTAGATGCCACGTATTTTTTCACGGAGC TGATAAAAATAAGAAGC |
| fecI-R3-R | CATAAAAATAAGAAGCCTGCAAATGCAGGCTTCTTATTTTTA TCAGCTCCGTGAAAAAA |
| rpoS-R1-F | GAATTTCTACTGTTGTAGATTCTTTTTCATCGGCCAGGATGTC CATAAAAATAAGAAGC |
| rpoS-R1-R | CATAAAAATAAGAAGCCTGCAAATGCAGGCTTCTTATTTTTA TGGACATCCTGGCCGAT |
| rpoS-R2-F | GAATTTCTACTGTTGTAGATAGCTCGAACAGCCATTTGACGA TGATAAAAATAAGAAGC |
| rpoS-R2-R | CATAAAAATAAGAAGCCTGCAAATGCAGGCTTCTTATTTTTA TCATCGTCAAATGGCTG |

| | |
|---|---|
| rpoD-R1-F | GAATTTCTACTGTTGTAGATATGGTCTCAATCATATGCACCGGAATAAAAATAAGAAGC |
| rpoD-R1-R | CATAAAAATAAGAAGCCTGCAAATGCAGGCTTCTTATTTTTATTCCGGTGCATATGATT |
| rpoD-R2-F | GAATTTCTACTGTTGTAGATATCATCACCGATCGGCGTTTCCATATAAAAATAAGAAGC |
| rpoD-R2-R | CATAAAAATAAGAAGCCTGCAAATGCAGGCTTCTTATTTTTATATGGAAACGCCGATCG |
| rpoD-R3-F | GAATTTCTACTGTTGTAGATGCGGGTAACGTCGAACTGTTTACCATAAAAATAAGAAGC |
| rpoD-R3-R | CATAAAAATAAGAAGCCTGCAAATGCAGGCTTCTTATTTTTATGGTAAACAGTTCGACG |
| rpoH-R1-F | GAATTTCTACTGTTGTAGATAATGCCGTCGGCAAAGTTAGATGAATAAAAATAAGAAGC |
| rpoH-R1-R | CATAAAAATAAGAAGCCTGCAAATGCAGGCTTCTTATTTTTATTCATCTAACTTTGCCG |
| rpoH-R2-F | GAATTTCTACTGTTGTAGATGTCCAGACCCTGCATCGCGTCGGTATAAAAATAAGAAGC |
| rpoH-R2-R | CATAAAAATAAGAAGCCTGCAAATGCAGGCTTCTTATTTTTATACCGACGCGATGCAGG |
| rpoH-R3-F | GAATTTCTACTGTTGTAGATCTGCAACGTGGACTTGTTGTCTTCATAAAAATAAGAAGC |
| rpoH-R3-R | CATAAAAATAAGAAGCCTGCAAATGCAGGCTTCTTATTTTTATGAAGACAACAAGTCCA |
| rpoE-R1-F | GAATTTCTACTGTTGTAGATTTGCCATGCGTAAATCTTCCGGGAATAAAAATAAGAAGC |
| rpoE-R1-R | CATAAAAATAAGAAGCCTGCAAATGCAGGCTTCTTATTTTTATTCCCGGAAGATTTACG |
| rpoE-R2-F | GAATTTCTACTGTTGTAGATATAGCTCAGGCCATCCAGCTCCCGATAAAAATAAGAAGC |
| rpoE-R2-R | CATAAAAATAAGAAGCCTGCAAATGCAGGCTTCTTATTTTTATCGGGAGCTGGATGGCC |

| | |
|---|---|
| rpoE-R3-F | GAATTTCTACTGTTGTAGATCCTCGCTCGGAAGATACGTGAACGATAAAAATAAGAAGC |
| rpoE-R3-R | CATAAAAATAAGAAGCCTGCAAATGCAGGCTTCTTATTTTTATCGTTCACGTATCTTCC |
| rpoN-R1-F | GAATTTCTACTGTTGTAGATCTGCTGTTCAACGATACAGCGACTATAAAAATAAGAAGC |
| rpoN-R1-R | CCATAAAAATAAGAAGCCTGCAAATGCAGGCTTCTTATTTTTATAGTCGCTGTATCGTT |
| rpoN-R2-F | GAATTTCTACTGTTGTAGATAGTTCAAAAATGCCTCGTGGACTAATAAAAATAAGAAGC |
| rpoN-R2-R | CATAAAAATAAGAAGCCTGCAAATGCAGGCTTCTTATTTTTATTAGTCCACGAGGCATT |
| rpoN-R3-F | GAATTTCTACTGTTGTAGATCTGTCGCTCAACGGTTTCGCTGGGATAAAAATAAGAAGC |
| rpoN-R3-R | CATAAAAATAAGAAGCCTGCAAATGCAGGCTTCTTATTTTTATCCCAGCGAAACCGTTG |
| filA-R1-F | GAATTTCTACTGTTGTAGATCTGTCCAGTAGTTGTTGTAGCGGGATAAAAATAAGAAGC |
| filA-R1-R | CATAAAAATAAGAAGCCTGCAAATGCAGGCTTCTTATTTTTATCCCGCTACAACAACTA |
| filA-R2-F | GAATTTCTACTGTTGTAGATGATGGCTTCCATCACCCGCTGGCGATAAAAATAAGAAGC |
| filA-R2-R | CATAAAAATAAGAAGCCTGCAAATGCAGGCTTCTTATTTTTATCGCCAGCGGGTGATGG |
| filA-R3-F | GAATTTCTACTGTTGTAGATATACCAGTTTTTCGCGCTCCGGCAATAAAAATAAGAAGC |
| filA-R3-R | CATAAAAATAAGAAGCCTGCAAATGCAGGCTTCTTATTTTTATTGCCGGAGCGCGAAAA |
| YfiQ-F | TTAGTTAAGTATAAGAAGGAGATATACATATGATGAGTCAGCGAGGACTGGAAG |
| YfiQ-R | GATGCTCATTTTGGCAGCGGCTTCACCGGAGCCTGATTCCTCGCGCTGGGCAAG |

| | |
|---|---|
| dCas (YfiQ)-F | GAGGAATCAGGCTCCGGTGAAGCCGCTGCCAAAATGAGCA TCTATCAGGAGTTC |
| dCas (YfiQ)-R | CCTCGATAAAGAATTGGTGGTACTTATCGATGATCTGCTTCG CTTTCTTATAG |
| RT-gapA-F | AACGTATCTGTAGTTGACCTGAC |
| RT-gapA-R | CAGGATACCAGTTTCACGAAG |
| RT-acs-F | ATTAAAGGTCAGGCGATCTACG |
| RT-acs-R | TATCGCCCAGGTTGCTGGTA |
| RT-rnb-F | ATGTTGGTGACTGGTTATACGC |
| RT-rnb-R | ATTTGTACAGTGCCGTTTTC |
| RT-adhE1-F | GGAGCCTTAAATGCAGGTATTG |
| RT-adhE1-R | CACCTGACCTAACCAGTTTATCAG |
| RT-rho-F | GTAAATGGAGAGAATCCAGAAAGGG |
| RT-rho-R | CTTTGGGCTATCTTTTTTAGAAGAGTAG |
| RT-gapA-2-F | GCATACATGCTGAAATATGACTCC |
| RT-gapA-2-R | TGATGTGTTTACGAGCAGTTTCG |
| RT-acs-2-F | CGGTAATGTGTCCATTAAATGGT |
| RT-acs-2-R | TCGAGCAGGGTATTGGCGAA |
| RT-rnb-2-F | AATCCGCAGAGCCAGAAGAAC |
| RT-rnb-2-R | CGATCGCCTTTCAGCGGAT |
| RT-rpoD-2-F | CATCTGGGGGATTTCATCGAGG |
| RT-rpoD-2-R | CTTCGCTTCGATCTGACGGATAC |

**Supplementary Table S3: The acetylated lysine residues of Acs with and without the TPA editor in vivo.**

| Protein | Sequences | Acetylated sites |
|---|---|---|
| Acs (At2-dCas12a) | K.VK*NTSFAPGNVSIK.W | 56K |
| | R.TAIIWEGDDASQSK*.H | 106K |
| | R.FANTLLELGIK*.K | 130K |
| | K.K*NVDDALK.N | 200K |
| | K.NVDDALK*NPNVTSVEHVVVLK.R | 207K |
| | K.NVDDALKNPNVTSVEHVVVLK*R.T | 221K |
| | R.MAQVVDK*HQVNILYTAPTAIR.A | 348K |
| | R.ALMAEGDK*AIEGTDR.S | 370K |
| | R.FEQTYFSTFK*.N | 493K |
| | R.LGTAEIESALVAHPK*.I | 541K |
| | R.SGK*IMR.R | 609K |
| | R.KIAAGDTSNLGDTSTLADPGVVEK*.L | 640K |
| Acs (dCas12a) | K.VK*NTSFAPGNVSIK.W | 56K |
| | R.TAIIWEGDDASQSK*.H | 106K |
| | R.FANTLLELGIK*.K | 130K |
| | K.K*GDVVAIYMPMVPEAAVAMLACAR.I | 131K |
| | K.NPNVTSVEHVVVLK*R.T | 161K |
| | K.K*NVDDALKNPNVTSVEHVVVLK.R | 200K |
| | K.NVDDALK*NPNVTSVEHVVVLK.R | 207K |
| | K.NVDDALKNPNVTSVEHVVVLK*R.T | 221K |
| | R.TGGK*IDWQEGR.D | 226K |
| | R.MAQVVDK*HQVNILYTAPTAIR.A | 348K |
| | R.ALMAEGDK*AIEGTDR.S | 370K |
| | R.FEQTYFSTFK*.N | 493K |
| | R.SGK*IMR.R | 609K |
| | R.K*IAAGDTSNLGDTSTLADPGVVEK.L | 617K |
| | R.KIAAGDTSNLGDTSTLADPGVVEK*.L | 640K |
